# A landscape of synergistic drug combinations in non-small-cell lung cancer

**DOI:** 10.1101/2021.06.03.447011

**Authors:** Nishanth Ulhas Nair, Patricia Greninger, Adam Friedman, Arnaud Amzallag, Eliane Cortez, Avinash Das Sahu, Joo Sang Lee, Anahita Dastur, Regina K. Egan, Ellen Murchie, Giovanna Stein Crowther, Joseph McClanaghan, Jessica Boisvert, Leah Damon, Jeffrey Ho, Angela Tam, Mathew J Garnett, Jeffrey A. Engelman, Daniel A. Haber, Eytan Ruppin, Cyril H. Benes

**Affiliations:** Cancer Data Science Laboratory, Center for Cancer Research, National Cancer Institute, National Institutes of Health, Bethesda, USA; Massachusetts General Hospital, Harvard Medical School, Boston, USA; Dana Farber Cancer Institute, Boston, USA; Samsung Medical Center, Sungkyunkwan University School of Medicine, Suwon 16419, Republic of Korea; Wellcome Trust Sanger Institute, Wellcome Trust Genome Campus, Cambridge, CB10 1SA, UK

## Abstract

Targeted therapeutics have advanced cancer treatment, but single agent activity remains limited by de novo and acquired resistance. Combining targeted drugs is broadly seen as a way to improve treatment outcome, motivating the ongoing search for efficacious combinations. To identify synergistic targeted therapy combinations and study the impact of tumor heterogeneity on combination outcome, we systematically tested over 5,000 two drug combinations at multiple doses across a collection of 81 non-small cancer cell lines. Both known and novel synergistic combinations were identified. Strikingly, very few combinations yield synergy across the majority of cell line models. Importantly, synergism mainly arises due to sensitization of single agent resistant models, rather than further sensitize already sensitive cell lines, frequently via dual targeting of a single or two highly interconnected pathways. This drug combinations resource, the largest of its kind should help delineate new synergistic regimens by facilitating the understanding of drug synergism in cancer.

## Introduction

Modern therapeutic approaches to numerous pathologies include the use of drug combinations to obtain better efficacy and lower systemic toxicity in patients. Combinations of drugs have been frequently used to treat microorganisms infections (Johnson and Perfect, 2010)(León-Buitimea et al., 2020), and most notably, tri-therapy against HIV infection can yield very long lasting disease control (Daar, 2017). Drug combinations are also frequently part of anti-cancer treatment, based mainly on empirical clinical discovery for decades (FREI et al., 1965) (Doroshow and Simon, 2017). Rationally designed targeted agents have now been approved across a variety of cancers but the vast majority of patients are still treated first with combinations of “classic” genotoxic chemotherapeutic agents such as DNA damaging agents or other agents targeting cycling cells (taxanes). Targeted agents are sometimes combined with traditional cytotoxics: e.g., the targeted agent trastuzumab (an antibody against HER2) is combined with taxane to achieve higher benefit in HER2 breast cancer (Marty et al., 2005) (Romond et al., 2005). Currently, there are only few combinations involving exclusively targeted agents that are used to treat cancer. There are however notable examples of recent successes: Combining CDK4/6 inhibition with Estrogen Receptor (ER) directed therapy is beneficial over other therapies in ER positive breast cancer (Schneeweiss et al., 2020). In AML, the BCL2 targeting agent venetoclax combined with the demethylating agent aza-cytidine provides substantial improvement in clinical outcome compared to either agent alone or chemotherapeutics regimens (Research, 2020). The use of BRAF and MEK1/2 inhibitors in combination has led to improved response in melanoma (Flaherty et al., 2012). Many other targeted combinations are now being tested in clinical trials.

While it stands to reason that combining targeted drugs could improve benefit, the rational development of drug combinations against cancer is still hampered by the limited understanding of underlying cellular processes. There is now ample evidence of heterogeneous response to targeted anti-cancer therapies even within molecularly stratified patients. Indeed, response is still highly variable within the best responsive patient cohorts, with treatment either inefficient up front (innate resistance) or of limited and unpredictable duration (acquired resistance) (Piotrowska et al., 2018). Whether combinations of targeted agents will show such heterogeneity in response or allow for more encompassing treatment regimen is not known. Another critical aspect, even for targeted agents, is toxicity. In contrast to drug combinations against HIV for example, targeted drugs against cancer address cellular processes that are almost always shared between cancer cells and normal cells. Consequently, even with targeted agents of good specificity, increased toxicity is a major hurdle for clinical development of combinations and is additionally very difficult to predict. To obtain higher efficacy than single agents and minimize systemic toxicity, drug combinations that are synergistic specifically in cancer cells are thus conceptually the most promising. Yet, the availability of public large scale combination datasets is limited, additionally impairing efficient computational modeling for combination discovery (Menden et al., 2019).

In this study we aimed to identify new combinations of interest that could help treat non-small-cell lung cancer (NSCLC) patients. Through a very large dataset we generated, we provide a robust estimate of the heterogeneity of response to targeted drug combinations within lung cancers and analyze genetic as well as cellular network determinants of synergism. This dataset will additionally provide a common grounds resource for the scientific community interested in drug combinations development against cancer, and in the development of computational modeling approaches towards the systematic discovery of synergism in cancer cells.

## Results

### A large-scale drug combination screen in NSCLC models, its design and scoring

To systematically study the response of non-small cell lung cancer models to pairwise drug combinations, a collection of 81 NSCLC cell lines that are genetically representative of human tumors (Garnett et al., 2012) was assembled. These models are extensively characterized at the molecular level (Iorio et al., 2016). Mutational profiles for major cancer genes in this collection are shown in **Supp Figure 1**. Similarly to what is seen in exome sequencing data of human tumors (Ghandi et al., 2019) only a handful of cancer genes (Tate et al., 2019) are recurrently mutated across the cell line collection (**Figure 1A**). Recently, fusion events were systematically identified for 79 out of 81 cell lines, most of which identified were not associated with a clear functional role (Picco et al., 2019), and thus were thus not studied for their relation to drug combination response here (except for EML4-ALK).

**Figure 1.**
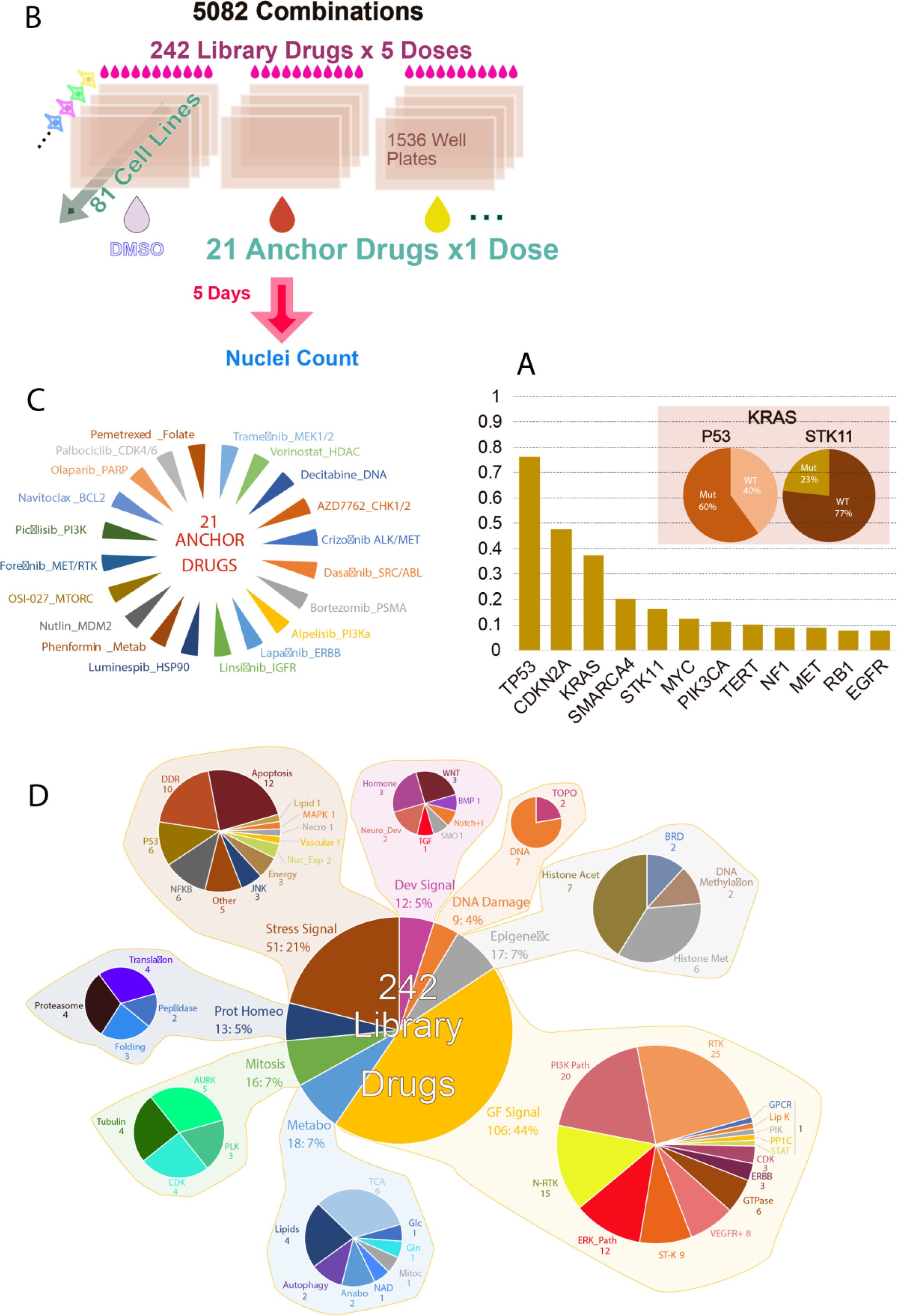
Overview of the study screening strategy. **A**. Major cancer drivers captured by the cell line collection screened. The y-axis in this figure shows the fraction of cell lines with a driver mutation. Figure also shows the percentage of KRAS mutated cell lines which also have a P53 and STK11 mutation. **B**. Screen set-up and key characteristics. **C**. Anchor drugs used. **D**. Library drugs used grouped by target class.

The response of the cell lines used in the present study to single-drug treatments was previously studied comprehensively across >400 single agents (Iorio et al., 2016). In addition, 49 of the cell lines were also part of a large chemical screening effort performed across NSCLC lines surveying an initial set of >200K compounds and an activity based selected subset of 447 chemical entities (Kim et al., 2013). These single-agent datasets as well as the results of genetic perturbations using shRNA (McDonald et al., 2017), super potent siRNA pools (Yuan et al., 2018) or more recently CRISPR CAS9 mediated loss of function (Tsherniak et al., 2017) (Behan et al., 2019) (Dempster et al., 2019) demonstrated that these NSCLC models capture the clinically relevant of therapeutic response of the disease. Importantly, as observed in the clinic, these data also demonstrate a prevalent heterogeneity of response to a given perturbation even within subsets of models sharing a common oncogenic driver (heterogeneity of response within KRAS driven NSCLC models for example, (Yuan et al., 2018)).

To identify synergistic drug pairs across the 81 cell lines, 21 “anchor” drugs were selected on the basis of their relevance to NSCLC treatment, approval status, results of preclinical therapeutic studies and biology. Those were combined with 242 “library” drugs covering the majority of targeted therapeutic classes currently in use or in development against cancer. This 21×242 testing strategy was used in an ultra-high throughput screen in 1536 well plates using one fixed dose of anchor drug and 5 doses for each library drug (**Figure 1B**, Supp. Tables S1, S2). **Figure 1C** lists the anchor drugs used and **Figure 1D** summarizes the targets and classes of library drugs. The dosing strategy of anchors and library drugs was aimed at discovering combinations with strong effect on viability (determined here using enumeration of nuclei across treatments). For this, drug dosages achieving complete or near complete targets suppression was sought. This strategy has previously been successful in discovering combinations to counter acquired resistance but is not conceptually restricted to this case (Crystal et al., 2014). The concentrations of the anchor drugs were chosen based on prior knowledge of on-target potency in cells and profile of response of these drugs across several hundreds of cell lines when available from prior studies (GDSC web site and unpublished data). A large-scale single-agent screen data was used to determine the dose of anchor drug that yielded very strong viability suppression in only a few cell lines (typically less than 2% of >500 cell lines tested). For EGFR inhibitors this would correspond to the highly sensitive cell lines that are dependent upon the EGFR mutant allele. The underlying concept is that while the outcome of target inhibition varies across cell lines, a given drug will overall affect its target(s) equivalently across cell lines (barring drug pumps effects which in fact do not strongly affect the vast majority of drug responses in cells (Iorio et al., 2016)). Thus, the anchor doses correspond to near complete suppression of target activity, which was for most targeted drugs ineffective in the majority of cell lines (Iorio et al., 2016). Similarly, for the library drugs, the concentration yielding strong viability suppression in only a few cell lines was determined based on single agent data or relevant literature. To further ensure that library drugs were suppressing their target(s) efficiently, one higher dose was added above this informed dose. Three additional lower doses were added to survey a larger breadth of target suppression. A dilution scheme of √10 was used (10-fold dilution every other dose). Drugs and concentration used are listed in Supp Table S1. The viability distribution for each single library drug and anchor across all doses is shown in **Supp Figure 1** demonstrating that the dosing strategy did yield an appropriately broad range of viability across cell lines.

The screen was performed in technical duplicates with two sets of identical plates seeded on a given day: two DMSO anchored plates corresponding to single agent treatments and two anchor plates corresponding to combination treatments. Screening was repeated for plates that failed quality control based on coefficient of variation (CV<25%) of the control wells (DMSO or anchor alone). To collect data on all anchors, a given cell line had to be seeded repeatedly on different days. With the goal of minimizing noise in the dataset, for each anchor, single agent testing (DMSO as anchor) was repeated in parallel with each anchor to allow matched DMSO anchored plates and combination plates of the same drugging run to be compared. Thus, throughout the analyses, combination (anchor plate) and single agent (DMSO plate) data are compared using only plates matched by cell seeding date. Prescreening calibration of the cell density allowing for proper proliferation and good cellular enumeration was performed. Failure rate varied across cell lines but was overall low: In total, 5,766 plates (1536 well plates) were used and 4,223 passed QC requiring the cell population to double at least once in addition to acceptable CV of control wells. 73 cell lines out of 81 tested had a pass rate above 90% and only two had a pass rate below 50%. Thus, while a small number of combinations were not captured in the QC passed dataset for a minority of cell lines, the overall data coverage is high, and the vast majority of tests were performed at least in two technical replicates (**Supp Figure 1**).

To evaluate the quality of the data the correlation of viability values across technical replicates was computed. There was an overall good correlation across technical replicates with Pearson’s R value across DMSO plates (single agent library + DMSO) of 0.80 and for technical replicates across combinations plates (Anchor drug + library drug) of 0.76. To evaluate the outcome and overall value of the screen data, a measure of synergy based on statistical independence of effect of the single agents was used (Bliss model): synergy was determined by considering the outcome of each single agent, taking the product of the single agent effects as the predicted outcome and comparing it to the experimentally determined viability outcome of the combination. The ratio between the expected and the observed outcomes constitutes the primary metric of synergy at each tested dose pair and an overall synergy score is derived from these 5 values (5 doses of library drug combined with one dose of anchor drug). A synergy score < 1 implies the drug combination is synergistic, with lower values indicating higher synergy (Methods, Supp. Note 1). To increase the likelihood of true positives we considered the 5 dose pairing individually and extracted the 2nd highest synergy from the series of 5 values, denoted as the *synergy score* of that combination. This 2nd best from the 5 synergy values can therefore correspond to any of the doses tested (not necessarily the 2nd maximum dose tested). To complement this synergy score an efficacy gain score was also computed: The *Higher than Single Agent (HSA) score* describes the additional viability loss observed with a combination over the maximum viability loss observed with either of its components individually. Here, a negative HSA score implies the drug combination is more effective than the better of the two drugs (Methods). To obtain a ranking of synergistic drug pairs, two complementary strategies were initially used, leveraging the synergy score: (i) computing the median synergy score across all tested cell lines and (ii) the count of cell lines with a synergy score of 0.8 or less (see below how different synergy score impact the number of synergies observed across cell lines). Similarly, for HSA, a global score was obtained by either (i) taking the median of all HSA scores for that combination across cell lines or (ii) counting the number of cell lines passing a threshold of 15% HSA (loss of 15% of cellular viability compared to lowest single agent viability).

Using these metrics, a set of combinations known to yield benefit over single agents or straightforwardly mechanistically supported were then scrutinized. For example, let us describe the results obtained with the anchor AZD7762 an inhibitor of the DNA damage repair response (DDR) kinases CHK1 and CHK2. Ranking combinations with AZD7762 based on the number of cell lines where the combination effect is either superior to single agents’ (HSA 15% or more) or synergistic (synergy score of 0.8 or less) shows that the inhibitor of Wee1, a kinase that regulates cell cycle checkpoint, is the top combination partner for AZD7762. Multiple synergies were also seen when combining ATR and CHK1/2 inhibitors (**Figure 2)**. There is published evidence for synergy between CHK1 and Wee1 inhibition (Guertin et al., 2012) (Aarts et al., 2015) (Buisson et al., 2015). The DNA damaging agents cytarabine, gemcitabine as well as the anti-metabolites pemetrexed and 5-FU also displayed HSA/Synergies in combination with AZD7762 albeit in fewer cell lines in the later cases than the former (**Figure 2**). Thus, there is clear detection of signal for combinations that were expected to be synergistic based on pathway knowledge and previous literature (Guertin et al., 2012; O’Neil et al., 2016). An overview of the screen outcome based on counts of synergy events across cell lines is presented in **Supp Figure 2** (see also Supp Table S2c).

**Figure 2.**
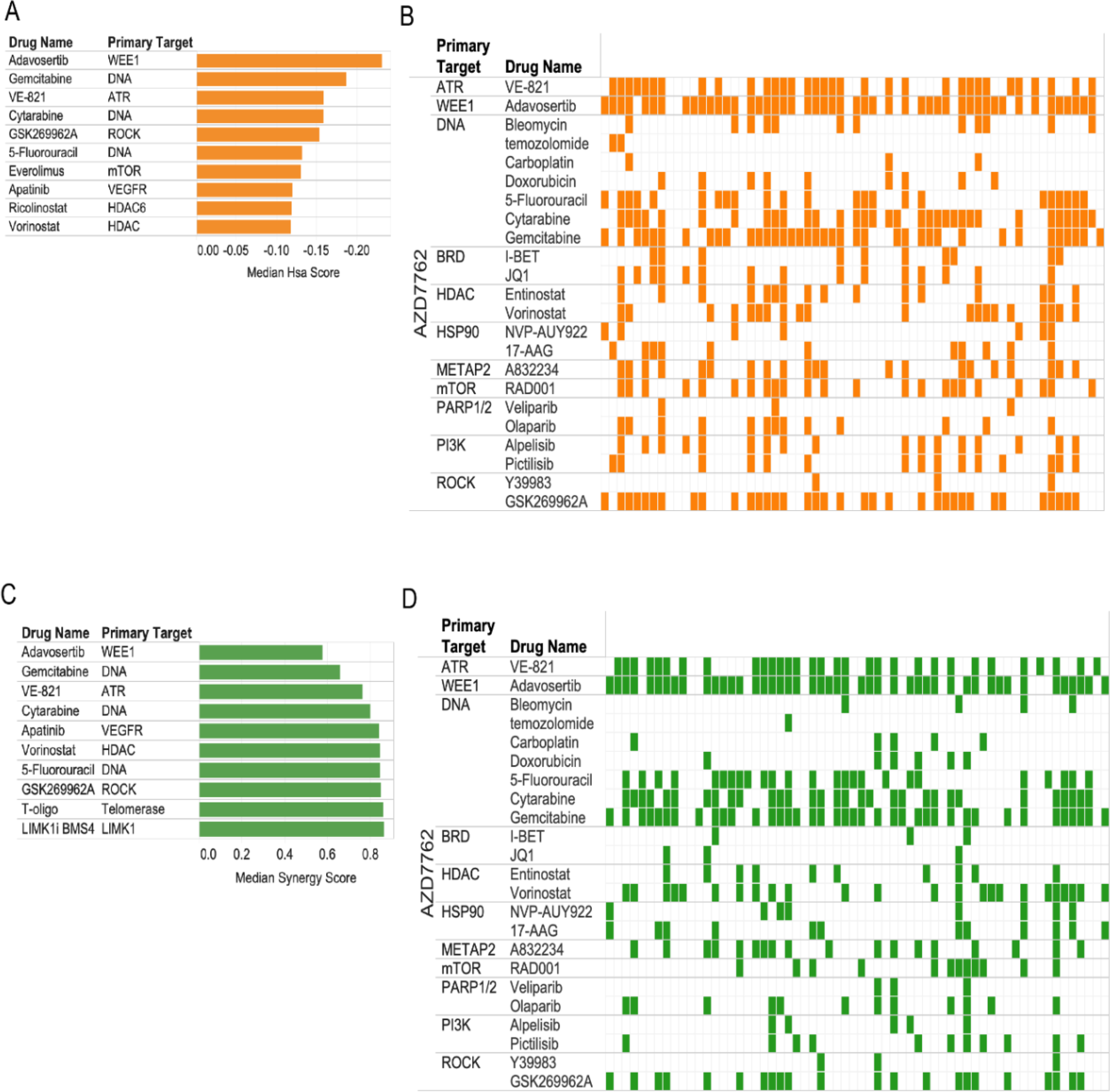
Capture of expected combinatorial effects. **A**. AZD7762 combinations: Top combinations sorted by median score across cell lines using Higher than Single Agent metric. **B**. Pattern of HSA events across top combinatorial partners for AZD7762 across cell lines. Each row corresponds to the specified drug combined with AZD7762 and each column (mark) corresponds to a cell line. Colored marks correspond to positive HSA events. The cell lines are in the same order across rows revealing differential pattern of HSA events for different combinations. **C**. Top combinations (with AZD7762) based on median synergy across cell lines. **D**. Pattern of synergistic events displayed as in B but using synergy rather than HSA.

To systematically identify top combinations for each anchor and estimate how impactful a given combination might be in the clinic, an *impact score* for each drug combination was computed based on the distribution of synergy scores across cell lines for each combination: This impact score was computed by comparing the distribution of synergy scores (or separately HSA scores) across cell lines for a given drug combination with the distribution of scores for all other drugs combined with the same anchor, using a Wilcoxon rank sum test. As a secondary measure, the median of the scores across cell lines for a given drug was compared to the median of scores of the rest of the drugs. The top combinations identified represent those with the highest effect across cell lines and thus perhaps across NSCLC patients. To further characterize the most promising combinations, the percent of cell lines with a synergy score within the top 5% of all scores (all anchors) was also computed. This systematic approach readily identified the combination of WEE1 inhibitor with CHK1/2 inhibitor and other combinations described above as the most impactful combinations for the CHK1/2 anchor (**Figure 3A**). Below we describe the top ranked combinations identified in our screens, based on their impact scores (**Figure 3G** shows a summary of the top combinations identified).

**Figure 3.**
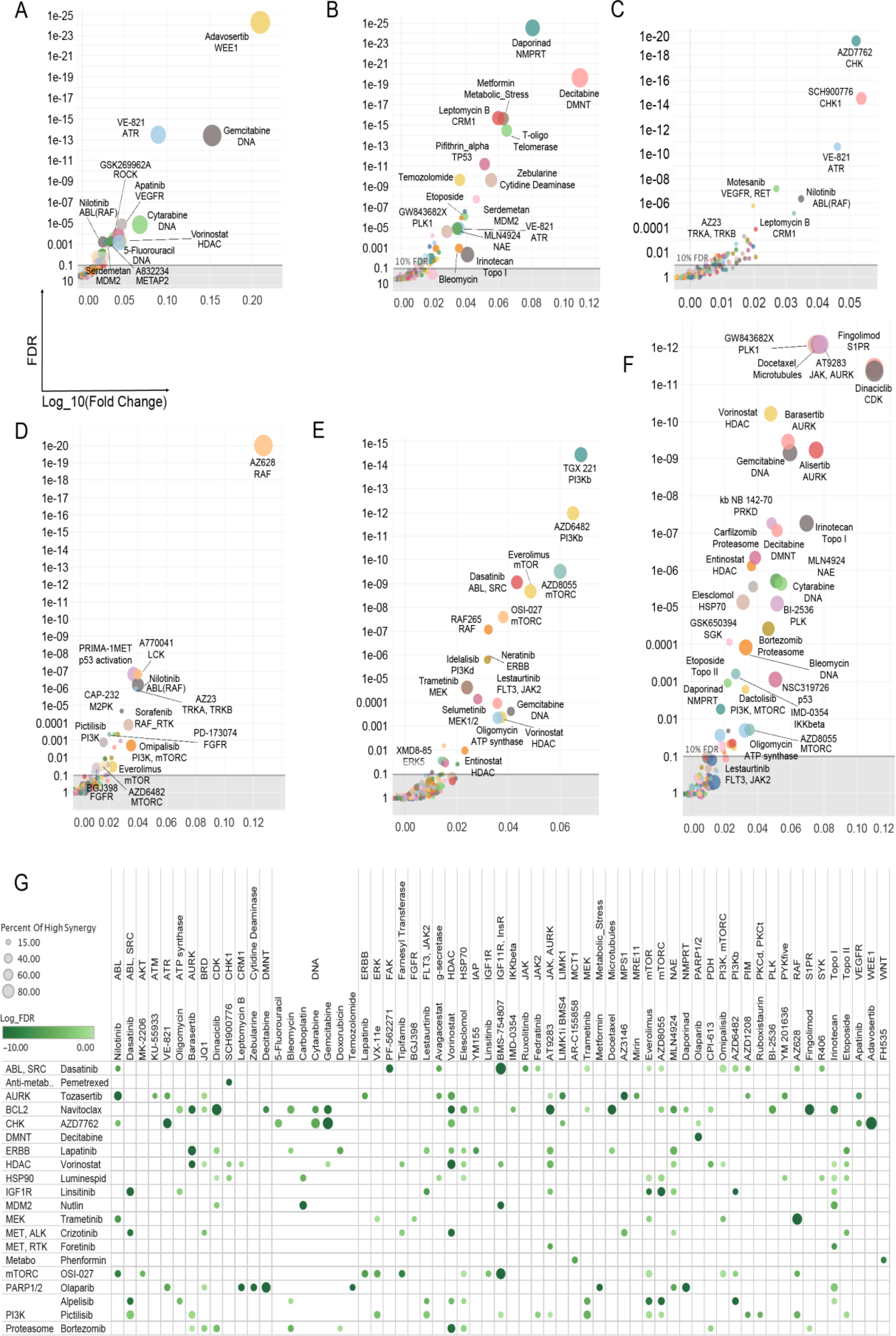
Top synergistic combination across anchors. **A-F**. The impact score for each combination is plotted with median synergy score of a given combination across cell lines compared to the median synergy score of all other pairs for that anchor represented on the X axis (as the Log10 ratio of median scores) and statistical enrichment of synergies for the plotted combination over all other tested combinations (with the same anchor) represented on the Y axis. The size of the dots represent the percentile of synergy scores for a given combination falling within the top 5% of all synergy scores for the whole screen (all anchors). Each plot corresponds to a different anchor drug: A, AZD7762 (CHK1/2); B, Olaparib (PARP); C, Pemetrexed (anti-folate); D, trametinib (MEK1/2); E, alpelisib (PI3Kalpha); F, navitoclax (BCL2). **G**. Overview of the top combinations based on impact score FDR and percentile of events (cell lines presenting with synergy) in the top 5% of all synergy scores (across all anchors: strong synergies). Combinations with at least 15% of strong synergistic events are shown (15% of the synergy scores across cell lines for that drug pair fall in the top 5% synergy scores overall). The size of the dots corresponds to percentile of events in the top 5% and color shade to the statistical enrichment (FDR).

### The landscape of the synergistic drug combinations uncovered

Here we review and discuss several interesting drug pairs synergistic across cell lines. We begin with describing cases that reaffirm known combinations that have been already reported in specific, dedicated studies, and move to new combinations that have not been yet reported in the literature.

We start with combinations involving the parylating enzymes Poly-ADP-Ribose-Polymerases (PARP), probably the most well-known examples of synthetic lethal based treatments to date. Somatic mutations BRCA1/2 are found across several cancer types and can confer clinical sensitivity to PARP inhibitors (Lord and Ashworth, 2017), such as olaparib. Top HSA partners we identified for olaparib include decitabine and zebularine, two related agents known to induce demethylation of DNA. Decitabine was also used as an anchor in the present study and olaparib was its top ranked synergistic partner with another PARP inhibitor veliparib ranking second (**Figure 3B**).

The MEK inhibitor trametinib (first approved by the FDA for use in BRAF V600E melanoma) has been studied in combination across a variety of contexts. Feedback re-activation of the MEK pathway upon suppression of MEK or ERK occurs impairs the clinical activity of BRAF inhibitors (Avraham and Yarden, 2011) (Lito et al., 2013) (Hatzivassiliou et al., 2013). Synergistic activity of RAF and MEK inhibitors combination has indeed been documented (Flobak et al., 2019) (Friedman et al., 2015) (Rukhlenko et al., 2018). Here, the pan RAF (A,B,C-RAF) inhibitor AZ628 was the top combination partner for trametinib. The ERK inhibitor VX11e also yielded frequent synergies with trametinib (**Figure 3G**). Consistent with published reports on treatment benefit in preclinical models (Engelman et al., 2008) (Alagesan et al., 2015) (Shapiro et al., 2019), synergies were also frequently observed between trametinib and inhibitors of the PI3K/mTOR pathway (**Figure 3D**, Supp Figure 2, 5C Across receptor tyrosine kinase inhibitors, those targeting insulin receptor / insulin growth factor receptor led to more synergies than combination with ERBB family members (**Supp Figure 3D**).

Drugs targeting the PI3K pathway also yielded interesting outcomes. The PI3K inhibitor alpelisib (BYL719) which targets selectively the alpha catalytic isoform of PI3K (encoded by PIK3CA, frequently mutated across multiple cancer types) was tested as an anchor drug. Combining BYL719 with PI3Kbeta selective inhibitors yields a strong HSA and synergistic effects across many cell lines with good consistency seen between the two PI3Kb inhibitors tested (AZD6482 and TGX221, **Figure 3E, Supp Figure 3-5**), in concordance with earlier reports in breast cancer (Costa et al., 2015). EGFR family inhibitors also display relatively frequent HSA with BYL719 across cell lines (Jänne et al., 2014) (Michmerhuizen et al., 2019). By contrast, the inhibition of other non-receptor tyrosine kinases of the SYK family or inhibition of FGFRs display no combinatorial benefit with BYL719. The pan-PI3K inhibitor pictilisib (GDC0941) and the MTORC inhibitor OSI-27 were also used as anchors: Pictilisib was broadly synergistic with trametinib, the ERK inhibitor VX11E and the mTOR inhibitor RAD001 (everolimus) (**Figure 3E, Supp Figure 3B**,**4**,**5C**). Similarly, pathway combinations of OSI-27 with PI3K and AKT inhibitors were synergistic in many cell lines, and ERK or MEK inhibitors were also among the top synergizing drugs with OSI027 (**Supp Figure 4-5, Figure 2**). Combining a catalytic inhibitor of MTORC1/2 and everolimus was previously shown to yield MTORC1 synergy (Nyfeler et al., 2012) and this was apparent here in the viability outcome. The insulin/insulin growth factor receptors inhibitor BMS754807 was the top RTK inhibitor synergizing with PI3K inhibition. Indeed, the insulin receptor family is a potent and major (even likely the ancestral) activator of PI3K amongst RTKs (Hopkins et al., 2018) (Hopkins et al., 2020).

CDK4/6 inhibition has been recently reported to be synthetic lethal with an array of partners. Here, the FDA approved CDK4/6 inhibitor palbociclib displayed strong synergies that match relatively well the recent data demonstrating the clinical relevance of the interaction between the inhibition of the PI3K/mTOR and CDK4/6 inhibition (Costa et al., 2019). MEK and ERK inhibition were also seen as producing some synergies with Palbociclib. Selicicilb (CDKs) and I-BET (BRD) were the top combination in terms of number of synergies. HSA analysis confirmed BRD targeting drugs JQ1 and I-BET as some of the top combinations with palbociclib (**Supp Figure 3**,**5**). The most cell lines with HSA were obtained with inhibition of mTOR and a number of strong HSA scores were seen with trametinib (de Leeuw et al., 2018) (Gopalan et al., 2018). Overall however, relatively few synergies were seen with Palbociclib and consequently their impact scores were low (which is why this anchor is not present in the overview presented in **Figure 3G**).

The inhibitors of the mitotic kinases AURK and PLK are among the drugs presenting with the most synergies with vorinostat, the anchor HDAC inhibitor. The proteasome inhibitor carfilzomib, the Nedd8 activating enzyme (NAE, involved in E3 Cullin family activation) inhibitor MLN4924, the BET inhibitors I-BET and JQ1, the LSD1 inhibitor LSD1-C76 and the topoisomerase I inhibitor irinotecan showed many synergies with vorinostat. These are well supported by literature in preclinical and for some, clinical studies (leukemia, cutaneous T-cell lymphoma for which vorinostat is an approved agent, multiple myeloma, (Suraweera et al., 2018). Synergies with all three AURK inhibitors tested are strong and numerous suggesting on-target basis for the observed effects. Synergy between one of the AURK inhibitors tested here, alisertib and the HDAC inhibitor romidepsin was reported in T-Cell lymphoma (Zullo et al., 2015). Agents targeting metabolic enzymes were also good combinatorial partners for vorinostat: CPI613 (PDH/aKGH), Bromopyruvate (Hexokinase) (**Supp Figure 4E**). Notably, we also find that AURK inhibitors had a clear tendency to be more broadly synergistic than CDK inhibitors (**Figure 4D, Supp Figure 7**).

**Figure 4:**
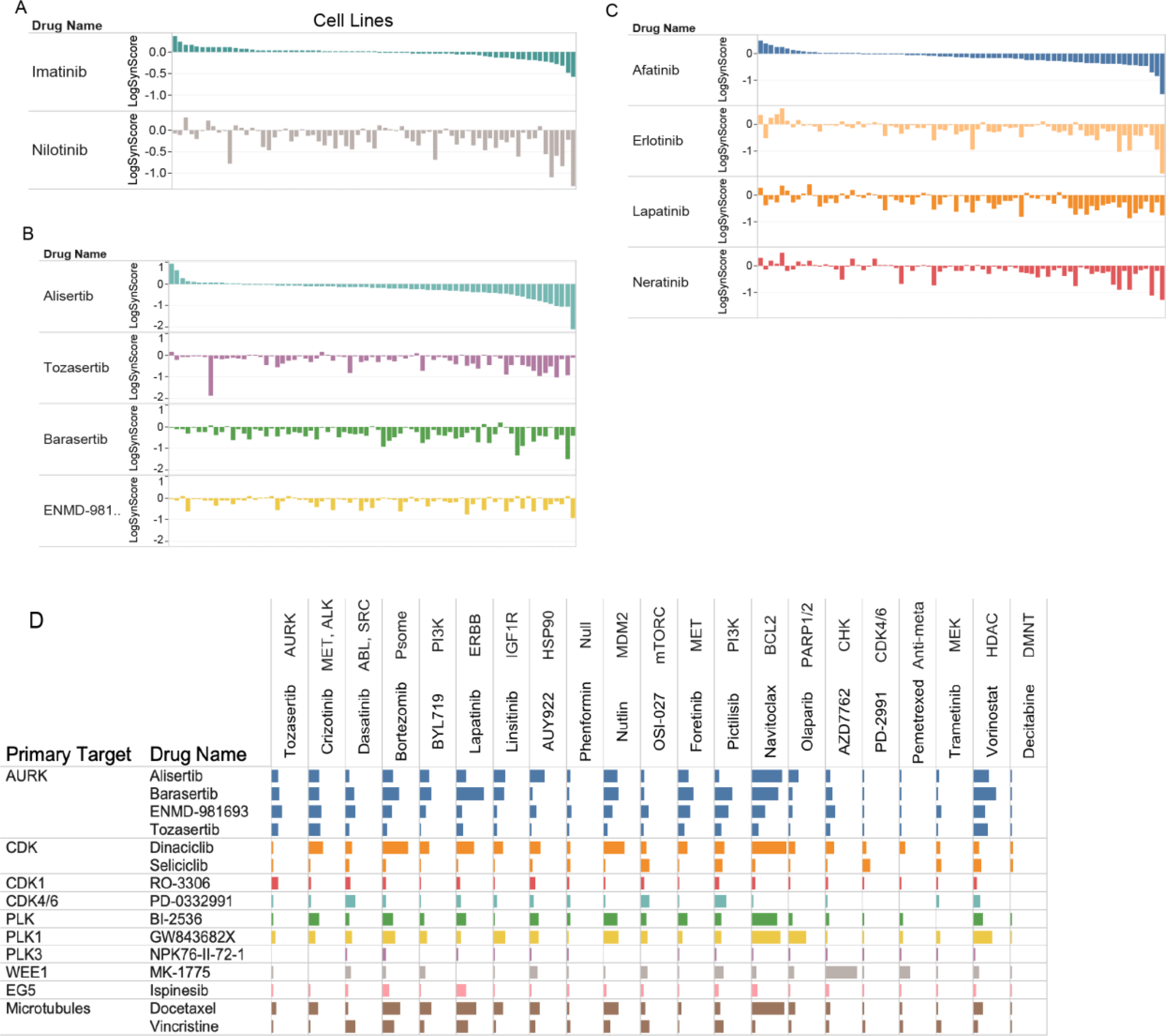
Differential synergistic outcome for mechanistically related drugs. **A-C**. The pattern of synergies for a given drug is presented as a ranked ordered plot of synergy scores for the first listed drug (top) compared to the synergy score of related drugs with cell lines in the same order as for the first drug. The log of the synergy score is plotted. **A:** Anchor drug is trametinib (MEK1/2); **B**. Anchor drug is vorinostat (HDAC); **C**. Anchor drug is OSI-027 (MTORC1/2). **D**. Overview of differential pattern for related library drugs across anchors. Selected library drugs with similar targets were chosen (rows) and the number of synergies in combination with the indicated anchors (columns) are plotted as bars (the bar size is proportional to percent of the synergy scores that are within the top 5% of all synergy scores).

Numerous growth factor pathway inhibitors synergize with the tyrosine kinase inhibitor dasatinib, an inhibitor of ABL, SRC family kinases (SFK) and multiple other tyrosine kinases. Multiple synergies were seen across inhibitors of the ERK and PI3K/mTOR pathways (**Supp Figure 3-5**). Consistent synergies were seen across EGFR family inhibitors and combination with INSR/IGF1R inhibitors yielded numerous HSA (**Supp Figure 4**). The FAK inhibitor PF-562271 synergized strongly with dasatinib across multiple cell lines, the JAK inhibitors TG101348 and ruxolitinib both synergize frequently with dasatinib albeit at a low level. Some synergies with the BTK family inhibitor ibrutinib (PCI32765, FDA approved for use against several hematological cancers which also inhibits BMX, a BTK family member expressed in carcinoma (Molina-Cerrillo et al., 2017) are also seen. All of these are in keeping with known signaling interactions between SFKs and FAK, BTK, JAK and RTK family members (Parsons and Parsons, 2004). Overall, dasatinib, perhaps due to its high level of polypharmacology is broadly synergistic but with top synergistic partners in keeping with known roles of SFKs in signal transduction.

A variety of other synergistic combinations emerged in the screen, which, due to space limitations are described in detail in the Supp. Note 3. Those include the finding that the BCL2 family inhibitor navitoclax is frequently synergistic with cell cycle blockers and strogly synergistic with the approved sphingosine receptor modulator fingolimod (**Figure 3F**), and the discovery of many additional synergistic combinations for which no previous report exists as far as we can tell, although in some instances indirect supporting evidence does exist. For example, the CHK1/2 inhibitor AZD7762 synergizes with the ROCK inhibitor GSK269962A (**Figure 2, 3A)** and a functional interaction between ROCK and DNA damage repair has been reported (Pranatharthi et al., 2019). AZD7762 also synergizes with the MetAP2 (Methionine Aminopeptidase) Inhibitor, A832234. There is precedent for regulation of cell cycle and MetAP targeting (Zhang et al., 2000).

Some of the strongest observed synergies are only found in very few cell lines and are thus not flagged by the impact score analysis presented in **Figures 2 and 3G**. Nevertheless, these highly context specific synergies might be mechanistically revealing and could be interesting to explore in other tumor types, where they might be more broadly relevant. The top synergistic combinations (top 5% of all synergy scores across all anchors) are represented as a network of Anchor-Library drugs interactions in Supp Figure 8. Interestingly, there is a high number of drugs that are shared between anchors among the top synergistic pairs. Perhaps pointing to core dependencies in the NSCLC lineage and to some biological processes and regions of the cellular interactome that could be prioritized for further explorations.

Finally, we note that although the present work focused on combinations of targeted agents, pemetrexed, a relatively well tolerated cytotoxic agent and one frequently used to treat NSCLC, was chosen as an anchor given that its administration is frequently associated with emerging resistance. A systematic screen of cytotoxic agents has recently been published across the NCI60 collection of cell lines (Holbeck et al., 2017). Consistent with its mechanism of action and previous studies (Grabauskiene et al., 2013), the top three synergistic drugs with pemetrexed were all inhibitors of the DNA damage response (**Figure 3C**). Few other strong synergies were detected across the rest of the library drugs including with motesanib (RTKs) and nilotinib (Abl, RAF, TKs). Interestingly, different cytotoxic agents gave distinct patterns of synergy even with drugs of similar MOA (such as DNA damaging agents). A striking example is the differential synergy profiles of vincristine and docetaxel. Both are targeting microtubules albeit through different mechanisms, but docetaxel displays many more synergies across anchors than vincristine. This doesn’t appear to be simply due to poor dosing choice for vincristine as there are instances of anchors displaying more synergies for vincristine than docetaxel (OSI027, Dasatinib, phenformin, **Figure 4D**). Overall, these results illustrate that there are likely important drug specific activities that need to be considered to select the most appropriate pairs of drugs (rather than only targets) (**Figure 3, Supp Figure 6**).

### Emerging properties of synergistic combinations discovered in our screen

Analysis of the characteristic properties of many of the synergistic combinations discovered reveals a few key emerging insights and principles, which we describe henceforth.

Because synergism emerges from the functional relation between targets there is considerable complexity to expect when drugs with multiple targets are combined. To study how polypharmacology (engagement of multiple often unrelated targets by a given drug) affects synergy, the synergy patterns of drugs sharing some targets but differing in others were compared. First, we begin with some notable cases. **Figure 4A** shows the outcome of the comparison between imatinib and nilotinib. Striking differences can be observed with a much larger number of strong synergies observed with nilotinib, which targets RAF in addition to ABL, which it targeted by both. Similarly, comparing four drugs targeting Aurora kinases (AURK) in combination with the HDAC inhibitor vorinostat (**Figure 4B**) and four different ERBB family inhibitors combined with the MTORC inhibitor OSI0927 (**Figure 4C**) shows that while the number of synergies and strength of those synergies are qualitatively similar, some synergies are unique to specific drugs. **Figure 4D** plots the synergy profile of library drugs targeting cell cycle entry and progression (see also **Supp Figure 3**). As expected, there is an overall similarity of their synergistic behavior across the majority of anchors. However, while genetic studies indicate some level of functional redundancy between CDK2,4 and 6, the independent targeting of either CDK2 (Dinaciclib) or CDK4/6 (Palbociclib, PD-03329921) can yield numerous different synergies. There are striking differences between CDKs inhibitors combinations with dinaciclib (Parry et al., 2010), showing a much more active profile than seliciclib (roscovitine, CDK1/2/5/9), with only few anchors including OSI-027 (mTORC) and palbociclib (CDK4/6) displaying more synergies with seliciclib than dinaciclib. Because both dinaciclib and seliciclib inhibit CDK1/2/5/9 equipotently (at least in vitro, (N’gompaza-Diarra et al., 2012)) it appears that secondary target(s) or perhaps differential mode of target engagement (Guiley et al., 2019) might be underlying the differences observed. Second, on a more general level, we find that combinations targeting just two targets (one single established target for each drug) are much less likely to be synergistic than combinations involving more than 2 targets (P=3.58×10^−34^ one-sided Wilcoxon test). There is also a mild but highly significant correlation between the total number of targets involved in a combination and percentile of cell lines in which it is synergistic (Spearman’s rho=0.21, P=6.25×10^−42^), (**Figure 5A**).Thus, drug specific effects are clearly seen and polypharmacology appears, as expected, to yield distinct context specific synergy outcomes, both at the drug and the cell-line levels. While this is likely to be an important hurdle for the rational development of combinatorial strategy, both in terms of efficacy and in terms of potential toxicities, it might also allow for the discovery of specific unexpected benefits (synergism) due to secondary target(s) inhibition.

**Figure 5:**
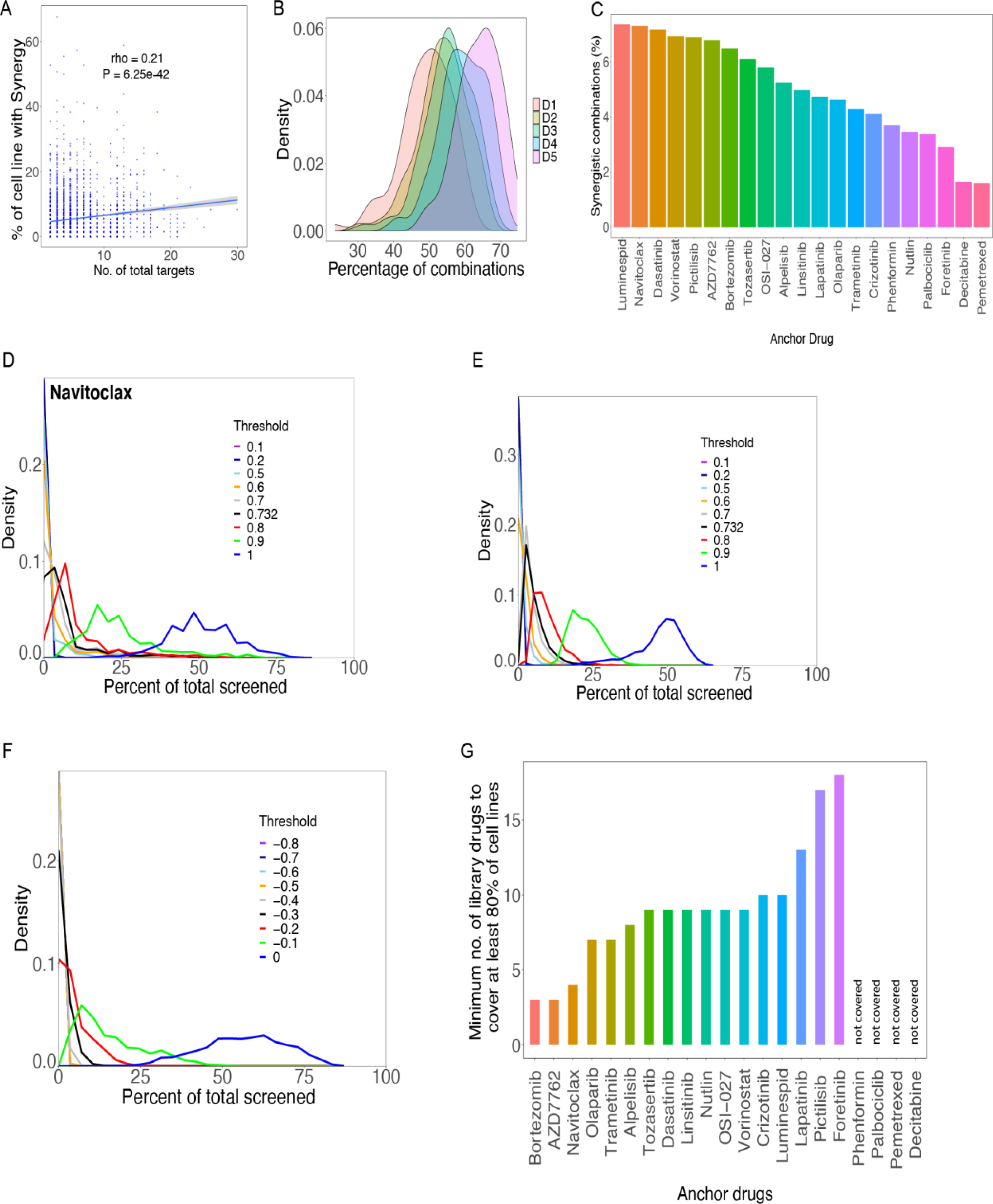
Sparsity of synergistic and HSA events across models. **A**. The number of synergistic events for a given combination depends on the number of targets of the combination. The percentage of cell lines presenting with synergy for a given combination is plotted against the number of targets addressed by the two drugs together (number of total targets). **B**. The proportion of cell lines harboring HSA increases with increased concentration of library drug. The density distribution of HSA events for each of the 5 doses of library drugs is plotted against the percent of cell lines presenting with HSA. **C**. Percent of synergistic combination for each anchor drugs as determined by the percent of cell lines presenting with a synergy score falling within the top 5% of all synergy scores. **D**. Density plots showing the fraction of synergistic events for anchor drug Navitoclax for various thresholds of synergy (see Methods for details). A low number of synergistic events is consistently observed using several synergy score thresholds. **E**. Same as D but across all anchors. **F**. Same as E but considered HSA events instead of synergistic events (Methods). **G**. Approximation of minimum number of library drugs needed to observe at least one synergistic event in at least 80% of the cell line collection.

A bird’s eye view of the results of our screen (**Supp Figure 2**) reveals that synergism typically occurs in a small number of cell lines and is thus strongly context dependent. Demonstrating that this sparsity of synergies is unlikely due to inappropriate dosing strategy, together with **Supp Figure 1, Figure 5B** shows that excessive dosing is not a likely broad cause of lack of HSA detection. Furthermore, the same sparsity property is seen with synergy scores. Because the synergy score is computed using a ratio of observed versus predicted outcomes, low viability outcome with single agents should not preclude detection of synergies. Consistent with this, there is only a very weak correlation between the viability outcome of the combination and the synergy score (Spearman rho= 0.038). To further study the sparsity of synergies across cell lines, the percentile of synergistic events for each anchor drug was computed defining strong synergy as the top 5% of all synergy scores observed in the screen. The HSP90 inhibitor Luminespid has the most synergies (∼7% of tests), followed closely by navitoclax. Pemetrexed and decitabine presented with the lowest number of synergies (below 2% of tests) (**Figure 5C**). The drug-drug network corresponding to the top 5% synergy scores can be found in Supp Figure 8. The coverage of cell lines (proportion of cell lines presenting with synergy) was computed for different thresholds of synergy, and remain always sparse (**Figure 5D**,**E**, Methods). A very similar pattern of distribution was observed when using a strong HSA score defined in an analogous manner (**Figure 5F**, Methods).

An important corollary of the high level of sparsity of synergy events observed is that multiple combinations would likely be needed to provide potentially effective treatments to a cohort of many different patients. To address this, we computed the number of library drugs that need to be combined with each anchor drug to obtain strong synergy in at least 80% of the cell lines. If these results would carry to the clinic, this would inform how many drugs might be considered to combine with an established agent in order to improve outcome for most patients. Notably, this analysis revealed four anchor drugs for which a coverage of 80% could not be achieved regardless of the number of combination partners used (see Supp. Note 2 for details). For the rest, the estimated number of drugs needed to obtain such coverage varied from 3 (bortezomib) to 18 (foretinib) (**Figure 5G**). Supp Table S3a contains results for different coverage thresholds. We note that a 100% coverage was obtained for only one anchor drug (Navitoclax, 14 drugs needed in combination).

We sought to understand whether adding a second drug tended to make sensitive cell lines furthermore sensitive or rather make already resistant cell lines sensitive (or both). Analysis of the effect of combinations (viability) in relation to the sensitivity observed with single agents revealed that the combination outcome is almost always contained within the range of sensitivity ever observed with single agents (Methods, **Figure 6A**). We term the rare drug pairs that diverge from this general pattern and actually yield an effect superior to what is seen with either agent alone in any cell-line as “super-sensitizers” (**Figure 6B**, Supp Tables S3b,c). Super-sensitizers are highly enriched in synergistic pairs (P=1.14×10^−25^, one-sided Wilcoxon test) (**Figure 6C**).

**Figure 6:**
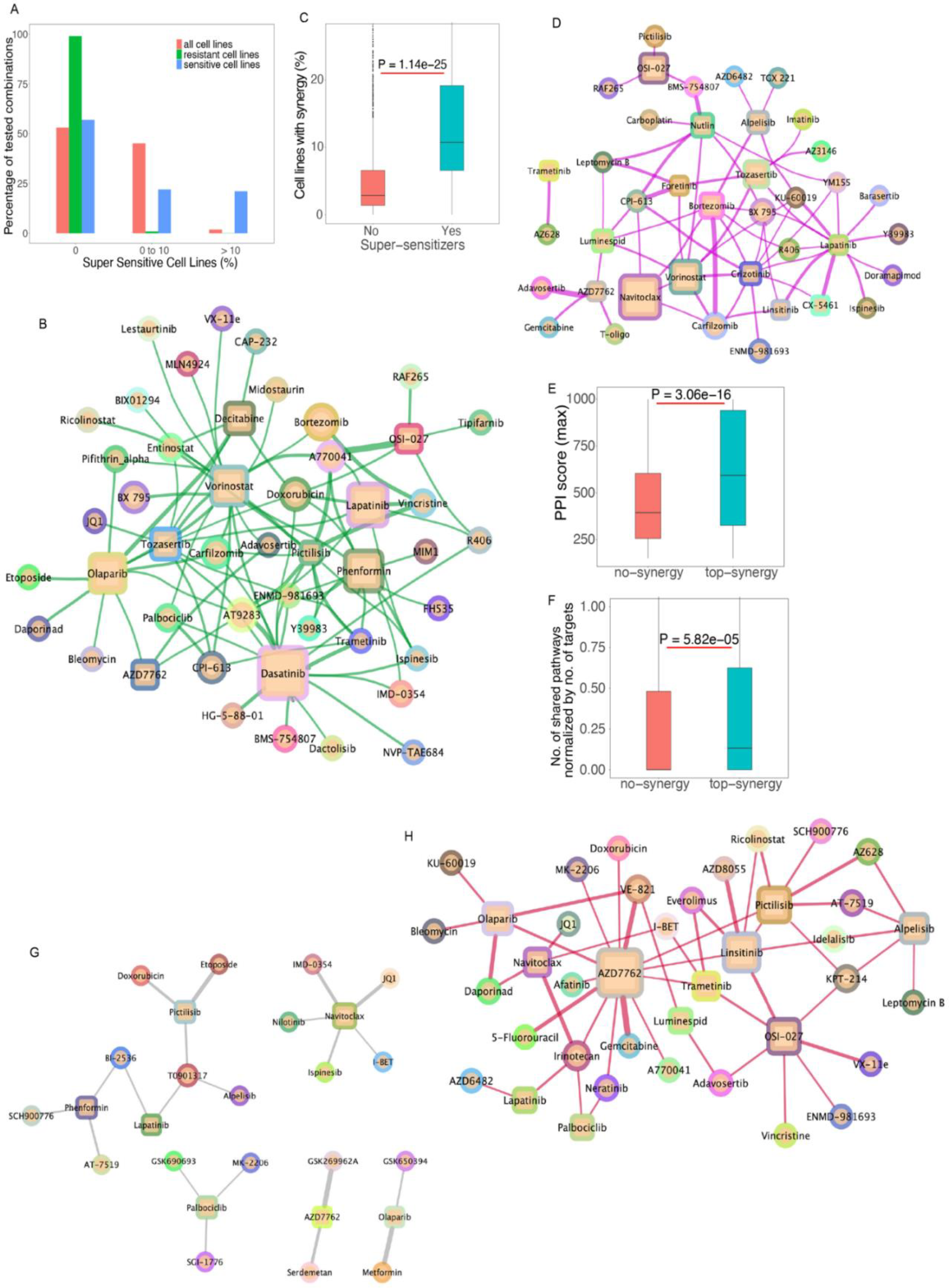
**A. Combinations rarely affect viability beyond effects observable with single agents.** The effect of combination treatment on each cell line was compared to the overall sensitivity to single agents observed across all cell lines. The percentile of cell lines in which the combination effect is superior to the effect of single agents in any cell lines (super sensitive cell lines) is shown in 3 categories (0, no observable supersensitive lines, 0 to 10 % and over 10%). The cell lines are further broken down based on their response to single agents (color). **B. Network view of drug combinations resulting in super-sensitization**. Anchors are displayed as squares and library drugs as circles. Library drugs that are also used as anchor drugs are represented by squares **C. Supersensitive events are enriched in synergies**. The percentile of synergistic events (cell lines) are compared between combinations that yield super sensitization versus those that do not. **D. Synthetic lethal drug pairs**: Network of drug combinations for drugs that yield substantial viability effect while single agents are deemed inactive. **E. Drugs with targets that are in close interaction with each other** based on previously determined protein-protein interaction network are more likely to yield synergies than those targeting un-connected proteins. **F. Drugs that target members of a given biological pathway** are more likely to yield synergy than those that target members of different pathways. G. **Synergistic drug pairs for which targets are encoded by genes engaged in a synthetic lethal interaction based on TCGA data analysis. H. Synergistic drug pairs with experimental evidence for a synthetic lethal relationship**.

Leveraging synthetic lethality has been a major focus of target discovery and therapeutic strategy development in oncology for some time (Kaelin, 2005) (Luo et al., 2009). In the context of drug combinations, we define synthetic lethal (SL) interactions when neither of single agents are markedly effective but the combination is (hence, these combinations correspond to a subset of more extreme synergistic interactions). To quantify these SL effects in our screen, inactive single agents (and doses) are designated as those yielding a viability greater than 75% of control treatment (DMSO), but the combination is strongly effective (defined as below 40% viability). The number of such instances is very low, comprising just 1.32% of all possible combinations (at library dose D4, 2nd max dose; **Figure 6D**). This result is reminiscent of the outcome of leveraging single agent sensitivity data in cell lines and combining single agent effective in a given set of cell lines to obtain synergy (Seashore-Ludlow et al., 2015). Interestingly, this result is comparable to fraction of synthetic lethal pairs seen in yeast and human cell line screens (Costanzo et al., 2016; Horlbeck et al., 2018; Srivas et al., 2016). While rare, these drug combinations are potentially of exceptional interest from a translational point of view (Supp table S3d). We further note that while true synthetic lethality (as strictly defined above) underlies a small minority of the synergistic combinations, evidently, by definition, synergism is essentially equivalent to synthetic sickness.

To study the impact of the cell-line cancer driver genotype on drug combination outcome, the percentage of highly synergistic combinations (top 5% synergy rank) across different genotypes was analyzed. Overall, across all anchors and drugs, there was no statistical imbalance for synergism for any of the major cancer driver genotypes, including KRAS, PIK3CA, EGFR, STK11. Interestingly, among KRAS mutant cell lines, STK11 (encoding LKB1) mutant cell lines were seen to harbor more synergies than STK11 WT ones. Although TP53 encodes a major tumor suppressor and sensor of cellular stress, TP53 mutations were not associated with synergism (or lack thereof). This is reminiscent of the results obtained with single agent treatment of very large cell line collections (Garnett et al., 2012)(Iorio et al., 2016). The analysis of the genetics of synergism for each anchor individually revealed that 3 anchors, crizotinib (MET / RTK), phenformin (ET Complex V) and AZD7762 (CHK1/2) were statistically more synergistic (number of top 5% synergies) in the KRAS mutant than in KRAS WT (one-sided Wilcoxon test, FDR<0.2). By contrast, 6 anchors promoted less synergies in STK11 mutant cell lines (KRAS WT and mutant) than STK11 WT ones: Tozasertib (VX-680, AURK), linsitinib (OSI-906, IGF1R), luminespid (AUY922, HSP90), phenformin (ETC), nutlin (MDM2), foretinib (XL-880, MET/ RTK). Additional results are presented in (Supp Table S4). When considering combination effectiveness rather than synergism, a number of combinations were seen to be more effective in a given mutant genotype (library dose D4, the second max dose was used for this analysis) (Supp Table S4): 195 for EGFR, 1128 for KRAS and 158 for PIK3CA mutated drivers (one-sided Wilcoxon test, FDR<0.2). As expected, most of the combinations showing higher effectiveness in EGFR mutant vs WT cell lines contained the EGFR inhibitor anchor lapatinib (144 out of 195). Similarly, for PIK3CA mutants, the PI3Kapha inhibitor BYL719 is present in all (158 out of 158) combinations showing higher effectiveness in PIK3CA compared to the WT. Interestingly, for KRAS, only 35 combinations involving trametinib showed an imbalance in effectiveness. Olaparib (AZD2281, PARP, 231 out of 1128), linsitinib (OSI-906, IGF1R, 150 out of 1128), navitoclax (ABT263, BCL2, 118 out of 1128) were anchors showing a high level of differential effectiveness in KRAS mutant versus WT models. Mutations in TP53 were associated with lower effectiveness for 199 combinations of which 184 involved nutlin (one-sided Wilcoxon test, FDR<0.2). This is expected since nutlin is predicably ineffective in TP53 mutant cell lines (Garnett et al., 2012). Contrary to expectations, we find that KRAS models did not respond to combinations treatment significantly differently from the KRAS WT cell-lines. This observation is further confirmed by a principal component analysis of the post-treatment viability values showing that KRAS WT and KRAS Mutant models do not segregate away from each other (**Supp Figure 9**).

Overall, the brunt of the synergistic events are hence not accounted for by the mutational state of recurrent cancer driver genes. This is in keeping with previous studies of drug combinations reporting a rather idiosyncratic pattern of synergy across models tested when classified on genotype alone (Horn et al., 2016) (Held et al., 2013) (Menden et al., 2019) (Friedman et al., 2015) (Flobak et al., 2019). Additionally, there was little difference in synergism or efficacy across subtypes of NSCLC (squamous, adenocarcinoma). Phenformin was the only anchor showing a subtype imbalance, with adenocarcinoma models harboring more synergies than squamous cell carcinoma models (P=0.00845, FDR<0.2, one-sided Wilcoxon test).

To gain a more general view of the characteristics of the targets involved in synergistic drug pairs, the protein-protein functional interaction (PPI) database STRING (Szklarczyk et al., 2019) was queried. This revealed that the targets of highly synergistic drug pairs are closer in the PPI network than the targets of non-synergistic drugs (P=3.06×10^−16^, one-sided Wilcoxon test) (**Figure 6E**). The same outcome was obtained using either all evidences or only experimentally validated PPIs in STRING and using median or maximum PPI score across targets. In an analogous manner, analysis of the KEGG pathway database (Kanehisa & Susumu, 2000; Carlson, 2016), showed that combinations targeting proteins within the same rather than different pathways are more likely to be synergistic (corrected for total number of targeted pathways to compensate for polypharmacology, see Methods, P=5.82×10^−05^, one-sided Wilcoxon test) (**Figure 6F**), in accordance with previous findings (Cheng et al., 2019) (Santolini and Barabási, 2018) (Lee et al., 2018) (Sahu et al., 2019).

To further evaluate the potential clinical relevance and benefit of the synergistic drug combinations identified here, a Cox regression analysis was performed using patient tumor data in TCGA. After controlling for single gene effect, age, gender, race, and cancer type among the 981 TCGA NSCLC, 43 drug combinations (6.04%), were seen to be linked via their targets downregulation to an improved patient survival (P < 0.05, Methods, Supp Table S5a). The BCL2-inhibitor navitoclax appears in 14 of these combinations. Repeating this analysis using copy-number data, showed that for 33 drug combinations low copy-number of the corresponding target pairs lead to improved predicted survival (Supp Table S5b). Here, the BCL2-inhibitor Navitoclax appeared in 17 combinations.

To uncover what fraction of the synergistic combinations identified in the screen might arise from SL interactions between their targets (this conceptually differs from the previous analysis presented above, where we quantified the direct SL-like interactions between the drugs themselves based on their phenotypic reduction of cell viability), we employed the ISLE pipeline (Lee et al., 2018) to analyze the lung cancer patient cohort of TCGA and identify the SL partners of each of the targets of the drugs screened in our analysis. Among the 1166 SL pairs found (Supp Table S6a), 83 matched to targets of tested combinations (Supp Table S6b). Notably, among those, 21 were synergistic (top 25% of synergy scores, Supp Table S6c, **Figure 6G**). Further, comparing the screened combinations to a published dataset of SL pairs identified in cell lines (Supp Table S6d; Ryan et al., 2018), showed that for 155 targets, at least one SL pair exists (Supp Table S6e) and that 55 of them are synergistic (**Figure 6H**, Supp. table S6f). Five drug combinations were common to both tumor derived and experimentally derived SL pairs, of which 2 were synergistic: Navitoclax + I-BET and Navitoclax + JQ1, that are mapping on the same targets (I-BET and JQ1 are both BRD targeting drugs). Thus, overall, SL analysis in patient data can explain a relatively small subset of synergistic combinations but might be useful to prioritize combinations found in the screens, indicating that they may be clinically relevant.

## Discussion

In this manuscript, we describe the outcome of a very large combinatorial drug screen surveying over 5,000 two drug combinations across 81 NSCLC highly characterized cell lines. By mining the literature on published drug combinations and using prior knowledge of cellular circuitry, we demonstrate the validity of both the data and the analytical strategy. Overall, we capture a large number of known or mechanistically transparent synergistic events that are consistent with prior knowledge. We also identify a considerable number of novel synergistic drug combinations. A subset of those have support from synthetic lethal analysis of NSCLC patient tumors data, which further testifies for their potential translational relevance. One of the most striking outcomes of our analyses is that synergistic combinations are mostly sparse and thus highly context specific. We find that combining together drugs that are not active as single agent almost never yields synergy. In addition, combining two drugs tends to render single agent resistant cell lines responsive rather than further sensitize already sensitive cell lines. Furthermore, sensitive cell lines rarely become super-sensitive, as combination effects mostly fall within the minimum viability levels observed for their individual components across all cell-lines. While this could potentially correspond to limited efficacy of combination of agents that are not efficacious on their own, exceptionally sensitizing combinations ca be found. In addition, synergism might more broadly provide benefit by allowing context specific activity of lower drug doses than used with single agents. The mutational status of major cancer genes is not highly predictive of synergy, as observed in other studies. Finally, synergy is more likely to emerge from targeting a single pathway or two interacting pathways, than by targeting two completely distinct pathways or functional modules of the cell. This finding is aligned with previous ones based on genetic perturbations in lower organisms (Cheng et al., 2019). One potential model explaining these findings is that when two combined drugs target sufficiently independent cellular functions then the highly evolved and robust homoeostatic control of the cellular system prevails. Thus, synergy, might often emerge from breaking homeostatic control.

There has been and still is considerable debate over what is synergy. Several competing models, that nevertheless often yield congruent conclusions (Tang et al., 2015) are used to qualify and quantify synergy. Here, a synergy scoring based on statistical independence akin to the broadly used Bliss model was used. We and others have previously demonstrated that this model is indeed valid to study viability outcome upon combinatorial treatment (Amzallag et al., 2019) (Flobak et al., 2019). Nevertheless, it is often pointed out that this type of modeling can in some instances assign synergy to cases of self-additivity. It is important to note that this counter intuitive outcome is limited to a small number of drugs. Examination of the relationship between self-synergy paradox and dose response shows that the drugs concerned have very steep dose response curves. Indeed, as seen here, vorinostat for example, has a much steeper dose response curve than most other drugs (see Supp. Note 4 for details). Our results indicate that synergy is overall a rare event, thus most drug combination are explained by the independent action of the two drug combined (as explicit in the Bliss hypothesis) which is aligned with recent modeling of clinical combination effectiveness (Palmer & Sorger, 2017). And their historical empirical development in cohorts of molecularly heterogeneous patients (Doroshow & Simon, 2017).

In summary, this work presents and analyzes the results of an exceptionally large dataset of drug combinations across lung cancer. The resulting dataset, that we make fully accessible to the scientific community, substantially expands on previous publicly available resources for drug combinations mining and modeling. There are many more analyses that could be performed using the data herein. Our hope is that these data will be subjected to additional analyses and foster the development of novel computational approaches towards a better understanding and prediction of drug-drug combinations outcome and the rules underlying synergistic interactions in cancer cells.

## Methods

### Drug Screening and Cell Viability Determination

Drug screening was performed using automated liquid handling in a 1536-well plate format.

The drug doses used were chosen based on previous single agent screening at the Center for Molecular Therapeutics of the Massachusetts General Hospital Center for Cancer Research.

The screen of two drug A and B was performed in a 1×5 format with 1 dose of drug A (anchor drug) combined to 5 doses of drug B (library drug) and compared to the effects of the 5 doses of drug B alone. The five doses of drug B followed a 4 fold dilution series. Screening was performed in replicate (two separate 1536 well plates).

Effect of drug treatment was determined by enumerating cell nuclei 5 days after the addition of drugs (day 0 designate the seeding day and day 1 the drug treatment day; no change of culture medium or drug re-addition were performed). Cells were seeded at densities optimized for proliferation based on pre-screen experimental determination in 1536 well plate format. Cells were seeded, placed overnight at 37oC and drugs added the next day using a pin tool. After 5 days in drug cells were fixed permeabilized and the cells’ nuclei stained in a single step by adding a PBS Triton X100 / Formaldehyde / Hoechst-33342 solution directly to the culture medium. Final concentrations: 0.05% TX-100 / 1% Formaldehyde / 1 ug/ml Hoechst-33342. Plates were covered and placed at 4oC until imaging. Imaging was performed on a ImageXpress Micro XL (Molecular Devices) using a 4X objective. Cell nuclei enumeration was performed using the MetaXpress software and count accuracy was routinely checked visually during acquisition.

Assay plates were simultaneously fixed (Para-Formaldehyde, 1% final concentration) and stained using the DNA intercalant Hoechst 33342 (final concentration 1 ug/ml) in the presence of 0.05% TX-100. No washing of plates was performed at any point post seeding. The plates were then imaged (4x magnification) on an automated microscope (ImageXpress Micro XL, Molecular Devices). The images captured the integrality of each well (1536 well plates) and nuclei were enumerated using the MetaXpress software (Molecular Devices).

Viability was computed as the ratio of number of nuclei in the drug treated wells over those in the control (DMSO treated) wells. For the drug combination plates, the anchor drug was added to all wells. The relative viability (compared to anchor alone treatment) was then computed by dividing the number of nuclei in treated (drug combination wells) by the anchor drug alone wells. This viability was then compared to the viability computed from the single agent wells (DMSO as an anchor). This allows for direct comparison of drug effect without using the values of the DMSO only wells (in the DMSO anchored plate) to compute drug combination effect. While mathematically equivalent to the cross plate comparison, this approach allows to minimize data noise due to potential plate to plate cell seeding number variation. Quality control criteria included a CV of less than 25% of the control wells (either DMSO alone wells for the DMSO anchored plates or Anchor alone in the Combination plates) and a cellular proliferation of at least 1 doubling. Proliferation was computed by comparing Day1 untreated plates (seeded concomitantly with the assay plates and fixed the day after seeding) to the DMSO only wells in the Day 6, DMSO anchor plates (Assay plates).

### Highest single agent (HSA) scores and synergy scores

We compute the HSA effect for each drug combination (on a cell line for a particular dose) by subtracting the combination viability minus the best (minimum) viability of the corresponding individual drugs. A negative value implies that the drug combination is more effective than the better of the two individual drug effects.

Synergy scores are computed using the Bliss model (Goldoni & Johansson, 2007; see Supp. Note 1 for details). The lower the score, the more synergistic the drug-combination is. A drug combination is synergistic if its score is less than 1. For each anchor-library-cell-line combination, we also compute the second-best synergy score among all the 5 library doses. We defined a drug combination to be highly synergistic in a cell line, if its synergy score (second-best, i.e. second-lowest) is less than a certain percentile (for example, 5 percentile or value of 0.732) of all combinations in all cell-lines. We show our results using various thresholds for detecting high-synergy combinations. We also compute the percentage of cell lines which are highly synergistic for each combination, and then rank all drug combinations based on this measure.

### Super-sensitizers

For each drug combination, we score the cell lines based on the max of the two drug effects (minimum viability). The 10 lowest ranked cell lines are considered to be resistant cell lines for the combination. The 10 top ranked cell lines are considered to be sensitive cell lines for the combination (excluding the most sensitive cell line). Now we check if the combination response (for library dose D4) in the resistant/sensitive cell lines is better than the best individual drug effect in the most sensitive cell line. If the combination effect is indeed better, we say that the combination makes the cell line super-sensitive. We considered all combinations which show super-sensitizer effect in at least 10 percent of the cell lines, and called them super-sensitizers.

### Drug-target mapping

We mapped the drugs to their targets using several resources: DrugBank (Wishart et al., 2017), Selleckchem.com. The mapping is shown in Supp Tables S1c,d.

### Protein-protein interaction (PPI) scores

We downloaded PPI network scores from the STRING database (Szklarczyk et al., 2019; downloaded on Aug. 8, 2019). We computed the PPI interaction score between the drug targets of any two drug combinations. If drugs have multiple targets, we either compute the max or median PPI score between the respective drug target pairs. We did his analysis using both the entire PPI network and by considering only drug target pairs which are bound to each other (called ‘binding’ in STRING).

The PPI score for the drug targets of the top synergistic combinations (based on top 5% synergy score overall) was computed based on all PPI information types in STRING. The PPI scores for the synergies with high scores in at least 10% of the cell lines tested (637 combinations) were compared to those for the non-synergistic pairs (lacking any strong synergy across cell lines, 743 combinations). For multi targeted drugs the maximum PPI score across targets was considered. This is for the analysis in Fig. 6E. The same results were obtained using only experimental binding evidence for PPI in STRING or using the median PPI score across targets rather than the maximum across targets.

### Patient survival analysis and synthetic lethal (SL) analysis

To test whether clinical survival benefit could potentially be derived from treatment with synergistic drug combinations identified, we mined data from 981 TCGA NSCLC patients (lung adenocarcinoma and lung squamous cell carcinoma patients). Only combinations whose targets could be mapped to TCGA gene set and with less than 4 targets per drug were considered because a search across a large number of targets might show a survival signal by chance. There were 712 such drug combinations among the top 25% most frequently synergistic combinations. For these combinations, low expression (below 33 percentile) of at least one of all potential target pairs yielded an improved survival benefit after controlling for single gene effect, age, gender, race, and cancer type among the 981 TCGA NSCLC patients (both lung adenocarcinoma and lung squamous cell carcinoma patients; Cancer Genome Atlas Research Network, 2012, 2014). The assumption being that the down-regulation of the target pair(s) may simulate clinical administration of the combination.

We used a computational method called ISLE (Lee *et al*., 2018) which mine 981 TCGA NSCLC patients (lung adenocarcinoma and lung squamous cell carcinoma patients) to identify clinically relevant SL pairs. ISLE uses 4 different filters: (a) It firsts mines *in vitro* shRNA/CRISPR datasets spanning hundreds of cell lines to identify potential SL candidates; (b) It then looks for negative selection of co-inactivated gene pairs using gene expression and copy number analysis in TCGA cancer patients; (c) ISLE then selects gene pairs whose co-inactivation (low expression or copy number) is associated with improved survival; (d) It finally selects SL pairs where the genes have high phylogenetic similarity. More details of this method is explained in (Lee *et al*., 2018). FDR threshold of 0.2 was used for this analysis. We mapped the drug combinations to their targets and mined for clinically relevant SL interactions between these target pairs. Only drug combinations where both the individual drugs have less than 4 targets are considered for this analysis. We identified experimentally derived SL gene pairs from various studies. The compilation of these various studies is provided in Cheng *et al*. (2021). There are 27975 experimentally identified SL pairs (Supp. Table S6d).

In both these analyses, a highly synergistic drug combination in a given cell line is by considering the top 5 percentile.

### Analysis for Figures 5D-F

For a fixed threshold of synergy or HSA, for each library drug at some dose, we look at all the anchor and cell line combination and check the fraction of them below the fixed synergy/HSA threshold (percentage of high HSA or high synergy). This will be our fraction of high synergies/HSAs for that library drug at some dose. We plot a density of these values for the fixed synergy/HSA threshold. We repeat the above procedure for different thresholds of synergy and HSA. We can also do the above procedure for a particular anchor drug.

## Acknowledgments

This research was supported in part by a grant from the Wellcome Trust (102696) and by the Intramural Research Program of the NIH, National Cancer Institute, and the Center for Cancer Research. This work used the computational resources of the NIH HPC Biowulf cluster (http://hpc.nih.gov). The results shown here are in part based upon data generated by the TCGA Research Network: https://www.cancer.gov/tcga. We thank Xeni Mitropoulos for her help on this manuscript.

