## Supplementary notes and supplementary figures for "A landscape of synergistic drug combinations in non-small-cell lung cancer"

### Supplementary notes and supplementary figures for paper "A landscape of synergistic drug combinations in non-small-cell lung cancer"

Nishanth Ulhas Nair[1], Patricia Greninger[2], Adam Friedman[2], Arnaud Amzallag[2], Eliane Cortez[2], Avinash Das Sahu[3], Joo Sang Lee[4], Anahita Dastur[2], Regina K. Egan[2], Ellen Murchie[2], Giovanna Stein Crowther[2], Joseph McClanaghan[2], Jessica Boisvert[2], Leah Damon[2], Jeffrey Ho[2], Angela Tam[2], Mathew J Garnett[2], Jeffrey A. Engelman[2], Daniel A. Haber[2], Eytan Ruppin[1,*], Cyril H. Benes[2,*]

1 – Cancer Data Science Laboratory, Center for Cancer Research, National Cancer Institute, National Institutes of Health, Bethesda, USA.
2 – Massachusetts General Hospital, Harvard Medical School, Boston, USA.
3 – Dana Farber Cancer Institute, Boston, USA
4 – Samsung Medical Center, Sungkyunkwan University School of Medicine, Suwon 16419, Republic of Korea.

#### Supplementary Notes

##### 1. Synergy computation

Synergy scores are computed using the Bliss model (Goldoni & Johansson, 2007). The lower the score, the more synergistic the drug-combination is. A drug combination is synergistic if its score is less than 1.

We defined a drug pair to be synergistic in the following manner: In each cell line and combination (for a particular library drug dose), we have cell counts measurements (day 6) upon library+anchor (LA) combination, library-only treatment (L), anchor-only treatment (A) and with only DMSO control treatment (C). Synergy score = (LA/A) / (L/C).

Consider the following example to calculate synergy:

| Cell Line | Anchor ID | Library ID | Cell Count (from drugged plates data) |
|---|---|---|---|
| X | DMSO | DMSO | 500 |
| X | DMSO | L | 450 |
| X | A | DMSO | 400 |

| | | | |
|---|---|---|---|
| X | A | L | 150 |

Library drug alone response Plate (I) compared to DMSO control = 450 / 500
Library + Anchor Plate (II) Compared to Anchor alone = 150 / 400

Synergy (delta from bliss) = (Library + Anchor Plate (II) Compared to Anchor alone) / Library drug alone response Plate (I) compared to DMSO control = (150/400) / (450/500)

We calculate synergy for each cell line and for every drug pair in each cell line for all doses D1 to D5 (for library drugs). We then compute the median value of the 2 replicates to get a synergy value for each drug pair for each cell line for a particular library dose.

**2. Coverage analysis**

We ask the question: for a given anchor, what is the minimum number of library drugs required so that at least 80% of the cell lines have one high synergy?

To address the above question, we consider the 21x242 drug combos across 81 cell lines. High synergy was defined based on second-best synergy (top 5% threshold). For each anchor, we have a binary matrix for all cell lines and libraries – a matrix of 1s and 0s, where 1 stands for high synergy and 0 otherwise. We used a simple greedy algorithm to compute the minimum no. of library drugs so that at least 80% of the cell lines have one high synergy. Since it is a greedy algorithm, it is an approximate solution.

The greedy algorithm used is similar to the classic greedy algorithm for the set covering problem (Chandu, 2015; Chvatal, 1979; Grossman & Wool, 1997).

Our greedy algorithm is as follows:

a) For a given anchor, we have a binary matrix (say M) for all cell lines and libraries – a matrix of 1s and 0s, where 1 stands for high synergy and 0 otherwise. Let D be the counter for the no. of drugs considered. Set D=0. Also set the cell lines considered as PC = NULL SET.

b) We pick the library drug with the maximum high synergy cell lines for the given anchor in matrix M. Set D = D + 1.

c) We remove those high synergy cell lines from M (let C be the names of those cell lines). So, we get a new smaller matrix for M. Calculate PC = union(PC, C).

d) Check if (length_of_PC / total_no_of_cell_lines)*100 < 80%. If so, repeat from step b.

We repeated the above analysis for different percentages of cell lines (from 50% to 100%).

##### 3. Potential mechanisms underlying some notable synergistic combinations

The first involves the NMPRT/NAMPT inhibitor daporinab (FK866), which was seen to yield HSA and synergies across many cell lines in combination with olaparib. PARP enzymes are thought to consume a large amount of cellular NAD as their substrate for PARylation on targets. Daporinab, by inhibiting NAMPT, lowers the levels of NAD in cells and this change in substrate availability has been previously suggested to underlie the synergy between PARP and NAMPT inhibitors (Bajrami et al., 2012) with potential impact for the treatment of Ewing's sarcoma (Heske et al., 2017).

Second, a number of synergies were observed with the insulin/insulin growth factor receptors inhibitor BMS754807. This outcome most likely corresponds to the well described feedback inhibition on IRS1(Insulin Receptor Substrate 1), a key adaptor in the Insulin Receptor pathway that is subject to inhibitory phosphorylation in an mTORC1 dependent manner and was recently described to imply degradation of IRS1 following phosphorylation by mTORC1 itself (Harrington et al., 2005; Yoneyama et al, 2018) BMS754807 synergizes less with BYL719 or GDC0941 than with OSI-027 even though mTORC1 is under the control of PI3K via AKT phosphorylation of TSC (Manning & Cantley, 2007). This is possibly due to the presence of other compensatory feedback loops that are not modulating the viability outcome when mTORC1 itself is targeted. In contrast to the PI3K inhibitors alpelisib (BYL719, PI3Kalpha) and pictilisib (GDC0941, pan PI3K), the MTORC inhibitor OSI027 synergizes broadly with the farnesyl transferase inhibitor tipifarnib (Figure 3G Figure 5-6). This is an unexpected outcome that seems to indicate either unappreciated activity of farnesyl transferase linked to mTORC1/2, perhaps linking HRAS to mTORC or an unsuspected off target of one of the two inhibitors. Interestingly, a recent study identified MTORC2 as a direct binding partner of RAS proteins albeit not specific to HRAS over other RAS isoforms (Kovalski et al., 2019).

Thirdly, targeting the RB pathway has been a long-standing interest in cancer therapeutics (Asghar et al., 2015). Inhibition of CDK4/6, which regulates cell cycle entry via the control of RB1/E2F complex, has in recent years been employed in multiple clinical settings. Single agent activity has been observed in preclinical models across different cancer types and promising results obtained in NSCLC models (Fry et al., 2004). In addition to improving the outcome of hormonal therapy in ER positive breast cancer inhibitors (Cristofanilli et al., 2016) of CDK4/6 are considered candidate combination agents in several cancers (Lim et al., 2016). CDK4 depletion in a mouse model of NSCLC driven by KRAS induces senescence and tumor regression (Puyol et al., 2010) and single agent activity of abemaciclib (LY2835219) a potent CDK4/6 inhibitor was seen in KRAS driven NSCLC human tumors (Patnaik et al., 2016). However, further exploration of this paradigm has not led to improvement in overall survival of patients in the single agent setting (Pacheco & Schenk, 2019). More recently, a preclinical study reported activity of CDK4/6 targeting in NSCLC with loss of SMARCA4 (a component of the SWI/SNF chromatin regulator complex; Xue et al., 2019). There is thus considerable interest in building combinatorial strategies around CDK4/6 inhibition, in particular for KRAS mutant NSCLC that have thus far not been successfully treated with targeted agents. Here we identify mTOR and MEK inhibitors as best combination partners for palbociclib (Supp Figure 3). However, overall, our results in vitro and with limited time of exposure to drugs, show a modest number of synergies with high sparsity across cell lines.

Nevertheless, longer time of treatment or non-cell autonomous effects could make these combinations beneficial in patients. Several clinical trials are poised to test the efficacy of MAPK pathway inhibitors in combination with CDK4/6 inhibition in NSCLC (for example, NCT03170206 for MAPK and NCT03065062 for mTOR/PI3K) and future combinations would be critical to address emerging resistance mechanisms (Wander et al., 2020). In addition, different CDK4/6 inhibitors vary in their target selectivity such that across the three approved CDK4/6 inhibitors, specific combinations might display differential synergies and/or clinical benefit (Hafner et al., 2019).

Our fourth set of interesting combinations involves SRC kinases. These enzymes, including p60c-Src and the closely related Src Family Kinases (SFK) members Fyn and Yes, are a group of non-receptor tyrosine kinases that have long been implicated in several hallmarks of cancer (Thomas & Brugge, 1997). In particular, SFKs regulate adhesion and motility mediated by integrins in coordination with the tyrosine kinase FAK as well as growth factor signaling. SFK are for example known to play a role in the activation of the MEK-ERK/MAPK pathway in several contexts and recent mechanistic studies support a model whereby SFKs can coordinate various signaling inputs with growth factor sensing by RTKs (Begley et al., 2015). Their targeting in tumors has long been elusive however and somewhat surprisingly, no mutations are found in these kinases in tumors. The multi-targeted kinase inhibitor dasatinib is a potent inhibitor of Bcr-Abl and was approved for use in Philadelphia positive chronic myelogenous leukemia (CML) and Acute Lymphoblastic Leukemia (Ph+ ALL) in 2006 (Jabbour & Kantarjian, 2020). It also potently inhibits Src and other SFKs and was used here to probe the potential of an approved drug targeting SFKs to yield beneficial combinations in NSCLC.  We found that TAE-684 synergizes broadly with dasatinib (Supp Figure 3,5). This is surprising since the kinase inhibitor TAE-684 was designed to inhibit the ALK tyrosine kinase which is an oncogene in NSCLC when fused to EML4 (Galkin et al). ALK is not normally expressed in NSCLC and ALK inhibitors are thus not broadly expected to be active outside of the ALK fusion driver context. This synergistic interaction between dasatinib and TAE-684 is thus most likely due to targeting of other kinase(s) than ALK, possibly IGF1R (Seashore-Ludlow et al., 2015). Two Insulin Receptor (IR) / Insulin Growth Factor 1 Receptor (IGF1R) inhibitors were used. Intriguingly, only one, BMS-754807 but not linsitinib displayed a very high number of synergies with dasatinib and indeed was the top synergizing drug with dasatinib. This strongly suggests that the synergies seen with the BMS compound are either not due to IR/IGF1R targeting or that additional targeting is needed to yield synergy (Supp Figure 4,5). This illustrates again the challenge brought by polypharmacology when trying to assign mechanisms to the observed synergies as well as how polypharmacology can result in differential synergistic outcome across compounds that are as single agents yielding similar activity profiles.

Fifth, a number of strong synergies were seen with the pan Aurora Kinase (AURK) inhibitor tozasertib (VX-680, used as an anchor) (Figure 3G, 4D,, Supp Figure 7-8). Three additional AURK inhibitors were used in the drug library: Barasertib (AZD-1152) which is selective for Aurora Kinase B over Aurora Kinase A, alisertib (MLN8237) and ENMD-981693 which are selective for Aurora kinase A. Synergistic activity was strongly detected between AURK inhibitors and HDAC inhibitors (Supp Figure 7) in line with previous reporting (Zullo et al., 2015). Interestingly, while both AURK and Cyclin Dependent Kinases are involved in cell cycle progression CDK inhibitors did not display the same broad pattern of synergies as AURK inhibitors (Supp Figure 7). Thus, cell cycle inhibition does not appear to be sufficient to explain the AURK inhibitors results and perhaps a more specific outcome of inhibiting mitotic progression with AURK inhibitors is at play or other non-mitotic targets (Bertolin et al., 2018; Otto et al., 2009; Bertolin & Tramier, 2020) of AURK are involved. Across anchors, different AURK inhibitors display differential synergy profiles. For example, while both barasertib and alisertib synergize with the BCL2 family targeting compound navitoclax, ENMD-981693 shows more synergies across cell lines with the CHK inhibitor AZD7762 than other AURK inhibitors. In contrast, all 4 AURK inhibitors tested synergize with the HDAC inhibitor vorinostat across many cell lines, albeit with only partial overlap of cell lines presenting synergy (Supp Figure 7). Thus, inhibition of either AURKA, AURKB (and AURKC) appears to have distinct outcomes in the combination setting. The mechanistic underlying of this observation is unclear but could be either a specific, possibly non-mitotic function of AURKs, a specific state of cell cycle arrest obtained with one inhibitor versus another, or alternate targets engaged by different inhibitors. Nevertheless, here and in other combination screens (Crystal et al., 2014) targeting AURK in combination with different growth factors, survival and DNA damaging agents could yield relatively frequent synergistic outcomes. Interestingly, AURKA has been linked to PI3K inhibition response in breast cancer and to resistance to EGFR inhibition in NSCLC (Donnella et al., 2018; Shah et al., 2019).

Sixth, the BCL2 family inhibitor synergizes with many drugs in the present study in keeping with previous results obtained in our studies on melanoma cell lines where navitoclax was characterized as a broad sensitizer to many other drugs (Friedman et al., 2015). This makes intuitive sense for a pro-apoptotic agent. In particular, some but not all inhibitors of the cell cycle present with numerous synergies when combined with navitoclax. For example, dinaciclib an inhibitor of multiple CDKs, BI-2536 and GW843682X inhibitors of Polo Like Kinases (PLK), or AT9283, alisertib and barasertib, inhibitors of AURKs are some of the drugs with the most synergies. There is precedent for the combination of dinaciclib and ABT263 to synergize via downregulation of MCL1, a major resistance factor to navitoclax as mentioned in the results section. Somewhat less expected are synergies observed between the ETC complex V inhibitor oligomycin and navitoclax, although a recent report on CLL identified Oxphos as a regulator or BCL2 inhibitor sensitivity and demonstrated synergy between oligomycin and BCL2 targeting in lymphoid cells (Guièze et al., 2019). BCL2 family members are not only involved in apoptosis by regulating the release of cytochrome C from the mitochondria but also in the maintenance of mitochondrial network function through regulation of fission-fusion cycle and possibly other means (Gross & Katz, 2017). Thus, the observed synergies between complex V inhibition and BCL2 inhibition could represent impact on mitochondrial integrity or sensitization to mitochondrial release of cytochrome C.

Seventh, the mitochondrial ATPsynthase inhibitor oligomycin was ranked 6th when considering median HSA score across cell lines for trametinib. This represented strong but relatively uncommon synergies and thus was also not flagged in the impact score analysis. Notably, oligomycin was the only drug with no obvious signaling network connection to MEK in the top 12 drugs ranked by median HSA. Oligomycin was also synergistic with OSI-27, pictilisib, dasatinib, navitoclax and perhaps less surprisingly with phenformin. These results suggest that disruption of ETC, while not necessarily lethal on its own, is creating vulnerability to core growth factor signaling inhibition by PI3K or ERK pathway inhibition.

#### 4. Self-addition breaking and limitation of statistical independence modeling

A simple explanation for self-additive coherence breaking is as follows: Consider a scenario where doubling the dose of a drug yields a doubling of the viability effect. Then consider what happens if the outcome of dose A for this drug is 80% viability. Doubling the dose (adding the same treatment to itself, Ax2), based on statistical independence would be expected to yield 64% viability (0.8x0.8). Thus, in this case yielding a synergy score of 0.625 (0.08/0.64). Now consider the same drug but with an initial dose yielding 40% viability. In this case the expected outcome is 16% (0.4x0.4; or 40% of 40%) and the observed outcome is 20%, meaning that no synergy is detected. Now, consider a drug that has a shallower dose response curve with a doubling of the dose yielding a 1.2-fold change in viability. This time, starting at 80% for dose A the expected outcome of 64% for (A)x2 is compared to an observed outcome of 0.67 (0.8/1.2) which does not represent synergy based on statistical independence (67% compared to 64%). Thus, both steepness of the curve and dosing (where on the dose response curve the data is acquired) impact the synergy call outcome. Importantly, analysis of single agent data in the present screen reveals that dosing yielded a broad range of viability values (Supp Figure 1) and that viability ratio for two consecutive doses chosen experimentally to be √10 apart (every other dose matches a 10-fold dilution) has a median value of 1.04 (1.14 for the top 25% ratio values). By contrast, the HDAC inhibitor vorinostat that shows self-additive synergy has a median ratio of viability across two consecutive doses of 1.44 (2.11 for the top 25% ratio values).

### Supplementary Figures

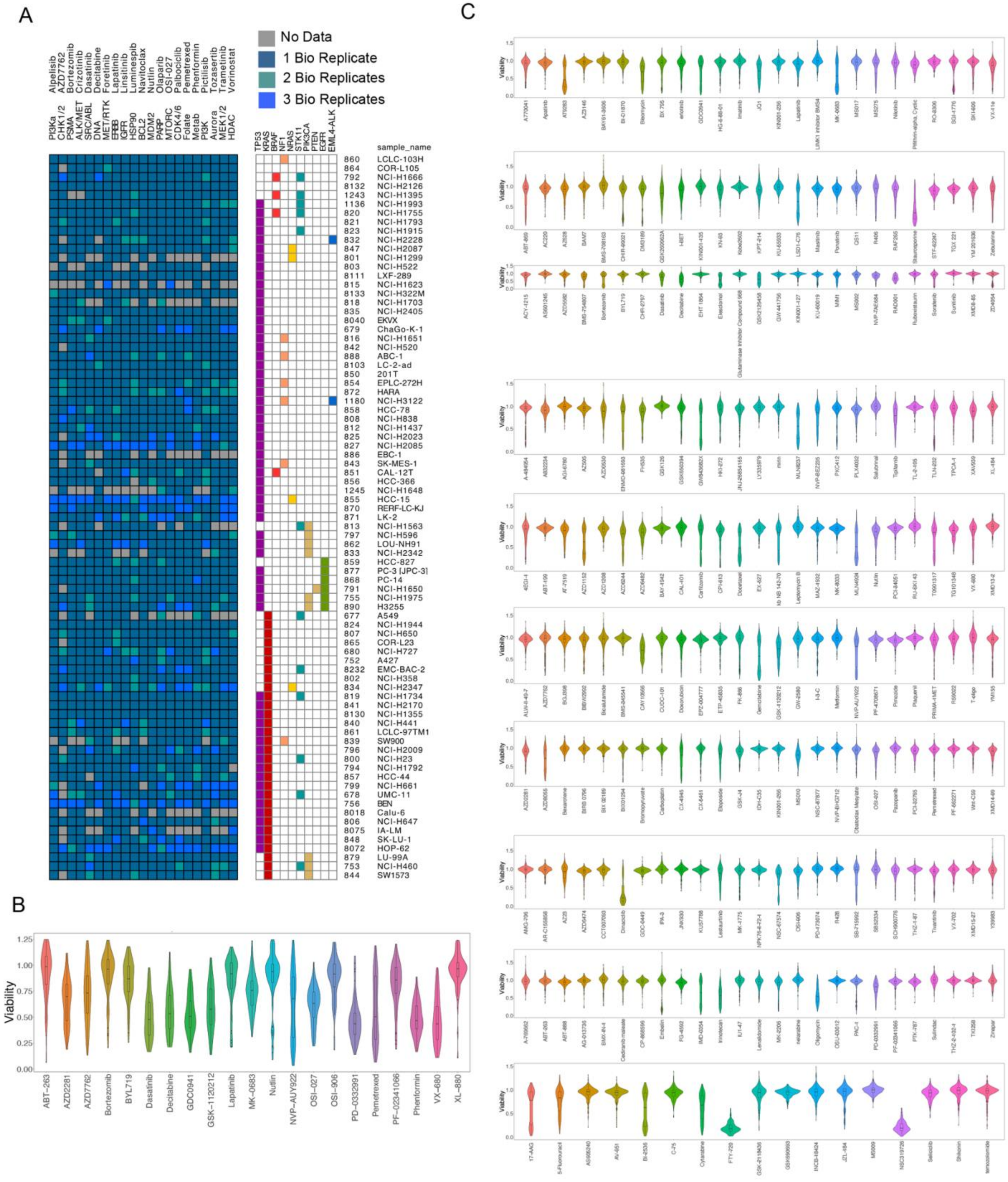
A
No Data
1 Bio Replicate
2 Bio Replicates
3 Bio Replicates
Alpelisib
AZD7762
Bortezomib
Crizotinib
Dasatinib
Decitabine
Foretinib
Lapatinib
Linsitinib
Luminespib
Navitoclax
Nutlin
Olaparib
OSI-027
Palbociclib
Pemetrexed
Phenformin
Pictilisib
Tozasertib
Trametinib
Vorinostat
PI3Ka
CHK1/2
PSMA
ALK/MET
SRC/ABL
DNA
MET/RTK
ERBB
IGFR
HSP90
BCL2
MDM2
PARP
MTORC
CDK4/6
Folate
Metab
PI3K
Aurora
MEK1/2
HDAC
TP53
KRAS
BRAF
NF1
NRAS
STK11
PIK3CA
PTEN
EGFR
EML4-ALK
sample_name
860 LCLC-103H
864 COR-L105
792 NCI-H1666
8132 NCI-H2126
1243 NCI-H1395
1136 NCI-H1993
820 NCI-H1755
821 NCI-H1793
823 NCI-H1915
832 NCI-H2228
847 NCI-H2087
801 NCI-H1299
803 NCI-H522
8111 LXF-289
815 NCI-H1623
8133 NCI-H322M
818 NCI-H1703
835 NCI-H2405
8040 EKVX
879 ChaGo-K-1
816 NCI-H1651
842 NCI-H520
888 ABC-1
8103 LC-2-ad
850 201T
854 EPLC-272H
872 HARA
1180 NCI-H3122
858 HCC-78
808 NCI-H838
812 NCI-H1437
825 NCI-H2023
827 NCI-H2085
886 EBC-1
843 SK-MES-1
851 CAL-12T
856 HCC-366
1245 NCI-H1648
855 HCC-15
870 RERF-LC-KJ
871 LK-2
813 NCI-H1563
797 NCI-H596
862 LOU-NH91
833 NCI-H2342
859 HCC-827
877 PC-3 [JPC-3]
868 PC-14
791 NCI-H1650
755 NCI-H1975
890 H3255
677 A549
824 NCI-H1944
807 NCI-H650
865 COR-L23
680 NCI-H727
752 A427
8232 EMC-BAC-2
802 NCI-H358
834 NCI-H2347
819 NCI-H1734
841 NCI-H2170
8130 NCI-H1355
840 NCI-H441
861 LCLC-97TM1
839 SW900
796 NCI-H2009
800 NCI-H23
794 NCI-H1792
857 HCC-44
799 NCI-H661
678 UMC-11
756 BEN
8018 Calu-6
806 NCI-H647
8075 IA-LM
848 SK-LU-1
8072 HOP-62
879 LU-99A
753 NCI-H460
844 SW1573
B
Viability
1.25
1.00
0.75
0.50
0.25
0.00
ABT-263
AZD2281
AZD7762
Bortezomib
BYL719
Dasatinib
Decitabine
GDC0941
GSK-1120212
Lapatinib
MK-0683
Nutlin
NVP-AUY922
OSI-027
OSI-906
PD-0332991
Pemetrexed
PF-02341066
Phenformin
VX-680
XL-880
C
Viability

**Supp Figure 1:** Overview of the dataset and response to single agents. A: Coverage of the cell line collection. The 81 cell lines (rows) are shown with key cancer genes altered to the right of the heatmap displaying data acquired and passing quality control. The number of biological replicates (Bio Replicates) corresponding to independent days of cell seeding are indicated by the color scheme indicated at the top right. B. Viability response to anchor drugs, the distribution of viability values (across all cell lines) is represented. C. Viability response to library drugs: For each library drugs the range of viability obtained with the indicated drug across the cell line collection is plotted. All doses are plotted together for each drug to demonstrate the overall range of viability obtained.

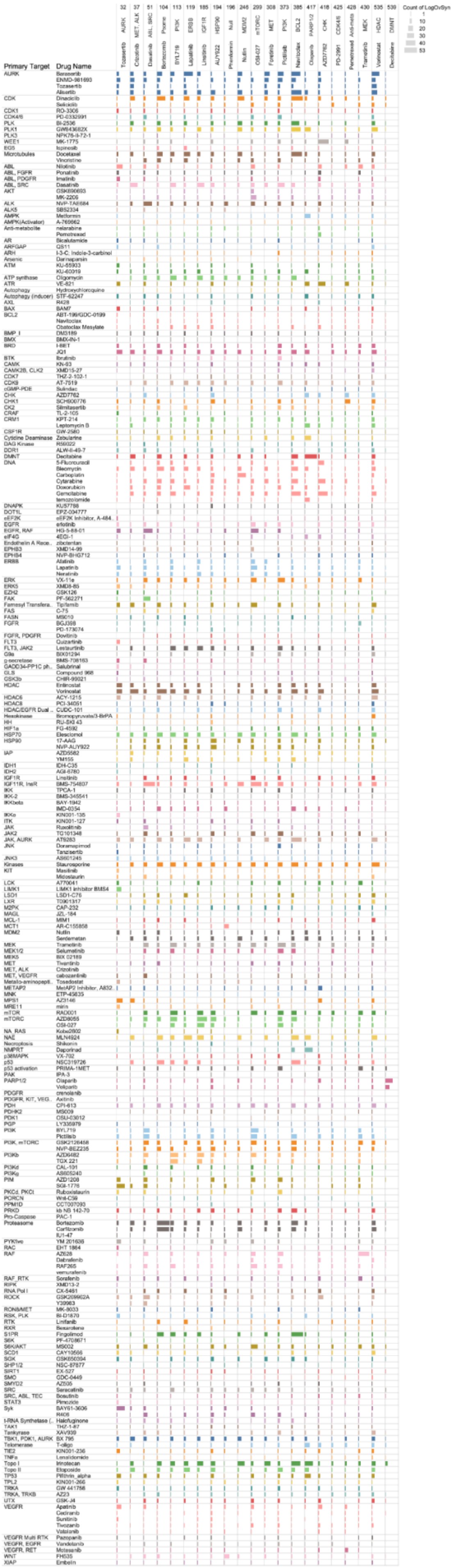

Primary Target
Drug Name
Count of LogOvSyn
1
10
20
30
40
53
32 AURK Tozasertib
37 MET, ALK Crizotinib
51 ABL, SRC Dasatinib
104 Psome Bortezomib
113 PI3K BYL719
119 ERBB Lapatinib
185 IGF1R Linsitinib
194 HSP90 AUY922
196 Null Phenformin
246 MDM2 Nutlin
299 mTORC OSI-027
308 MET Foretinib
373 PI3K Pictilisib
385 BCL2 Navitoclax
417 PARP1/2 Olaparib
418 CHK AZD7762
425 CDK4/6 PD-2991
428 Anti-mets Pemetrexed
430 MEK Trametinib
535 HDAC Vorinostat
539 DMNT Decitabine

**Supp Figure 2:** Profile of synergy counts across all tested combinations. For each combination between the indicated library drug (row) and anchor drug (column) the number of cell lines harboring synergy is depicted by a bar. The size of the bar is proportional to the number of cell lines harboring synergy with maximum size set at 53 (maximum number of cell lines presenting with synergy). Synergy count is based on a synergy score threshold of 0.8.

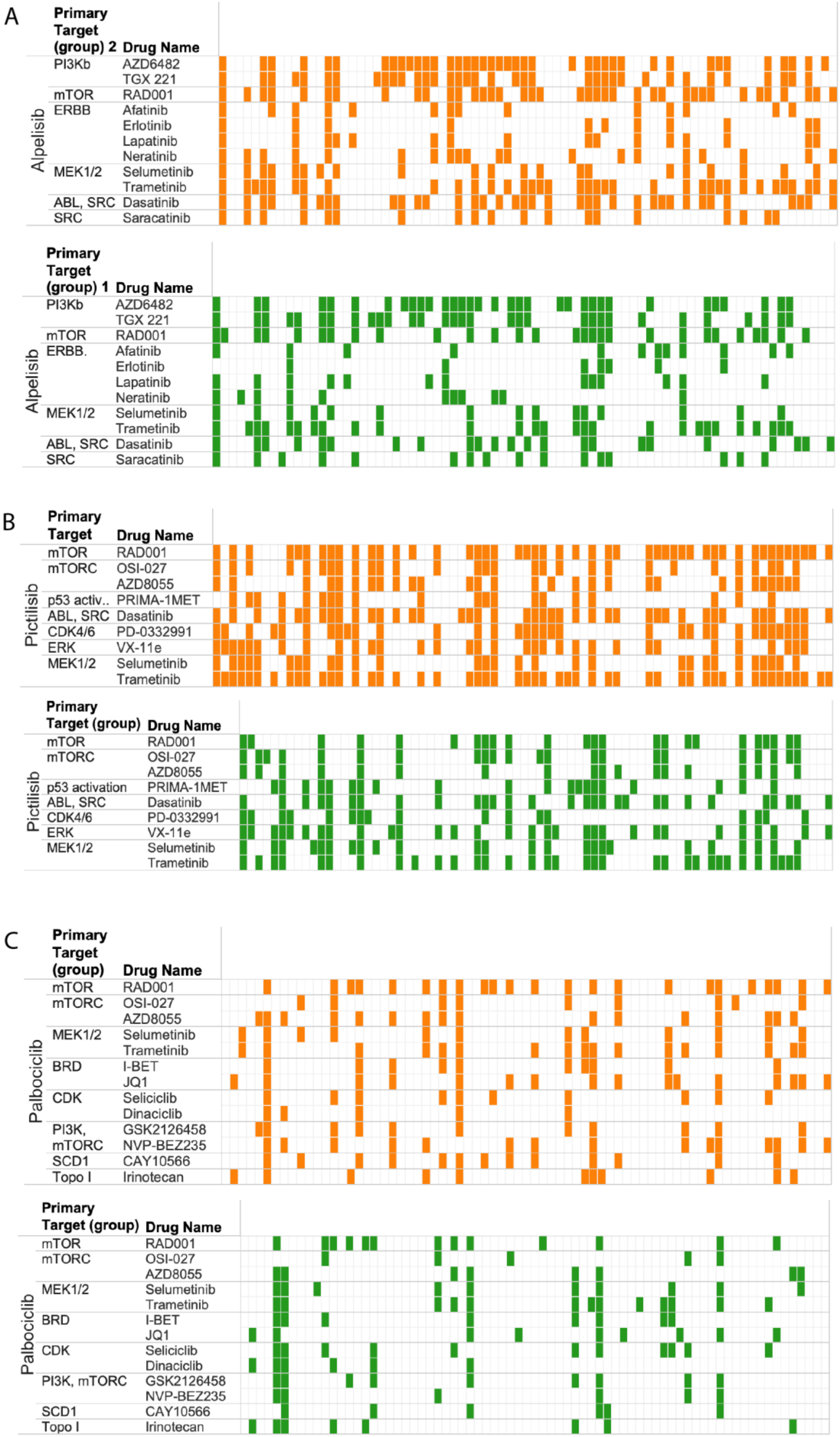
A
Primary Target (group) 2
Drug Name
Alpelisib
PI3Kb AZD6482
TGX 221
mTOR RAD001
ERBB Afatinib
Erlotinib
Lapatinib
Neratinib
MEK1/2 Selumetinib
Trametinib
ABL, SRC Dasatinib
SRC Saracatinib
Primary Target (group) 1
Drug Name
Alpelisib
PI3Kb AZD6482
TGX 221
mTOR RAD001
ERBB. Afatinib
Erlotinib
Lapatinib
Neratinib
MEK1/2 Selumetinib
Trametinib
ABL, SRC Dasatinib
SRC Saracatinib
B
Primary Target
Drug Name
Pictilisib
mTOR RAD001
mTORC OSI-027
AZD8055
p53 activ.. PRIMA-1MET
ABL, SRC Dasatinib
CDK4/6 PD-0332991
ERK VX-11e
MEK1/2 Selumetinib
Trametinib
Primary Target (group)
Drug Name
Pictilisib
mTOR RAD001
mTORC OSI-027
AZD8055
p53 activation PRIMA-1MET
ABL, SRC Dasatinib
CDK4/6 PD-0332991
ERK VX-11e
MEK1/2 Selumetinib
Trametinib
C
Primary Target (group)
Drug Name
Palbociclib
mTOR RAD001
mTORC OSI-027
AZD8055
MEK1/2 Selumetinib
Trametinib
BRD I-BET
JQ1
CDK Seliciclib
Dinaciclib
PI3K, mTORC GSK2126458
NVP-BEZ235
SCD1 CAY10566
Topo I Irinotecan
Primary Target (group)
Drug Name
Palbociclib
mTOR RAD001
mTORC OSI-027
AZD8055
MEK1/2 Selumetinib
Trametinib
BRD I-BET
JQ1
CDK Seliciclib
Dinaciclib
PI3K, mTORC GSK2126458
NVP-BEZ235
SCD1 CAY10566
Topo I Irinotecan

D

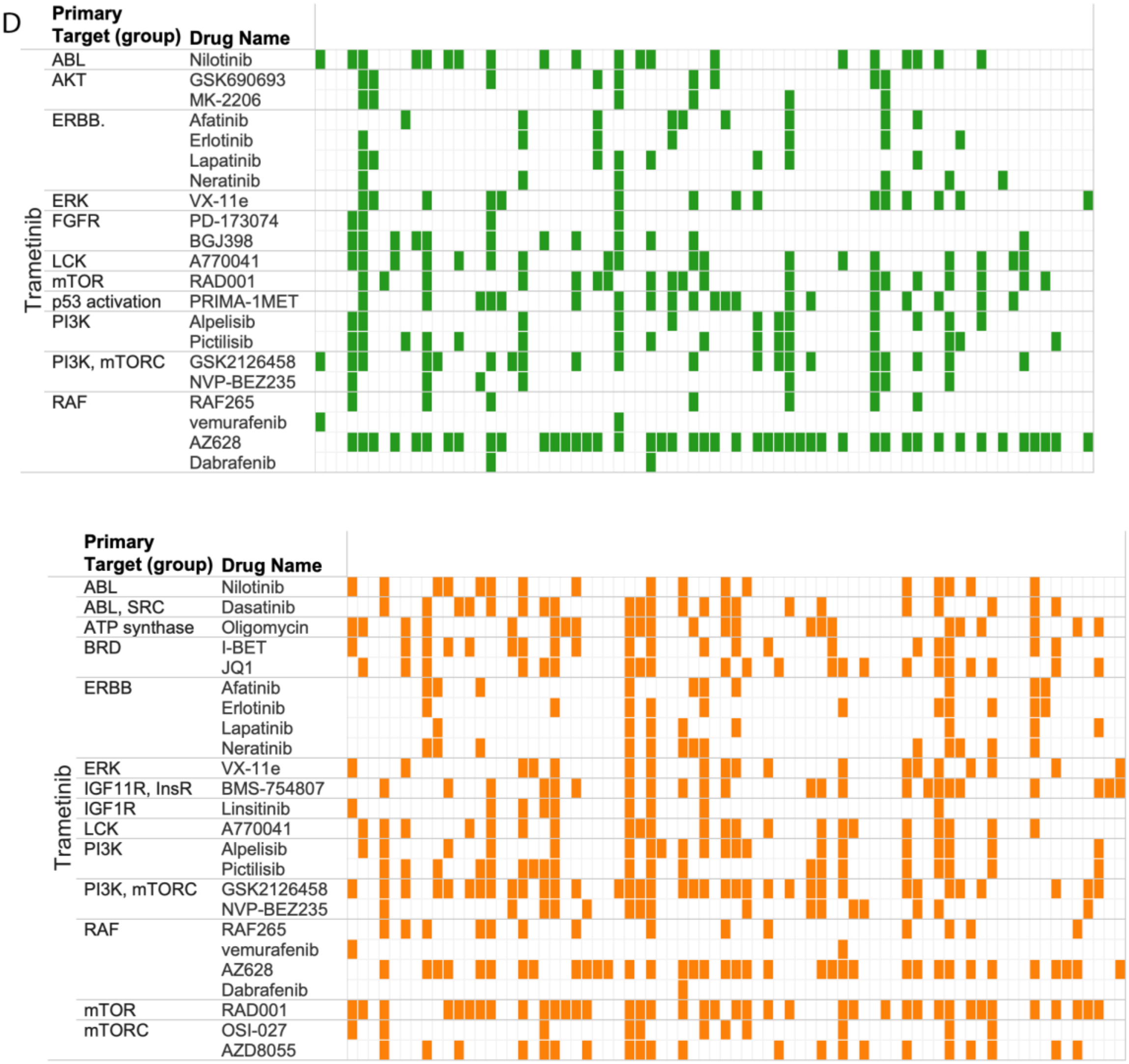

Primary Target (group)
Drug Name
Trametinib
ABL Nilotinib
AKT GSK690693
MK-2206
ERBB. Afatinib
Erlotinib
Lapatinib
Neratinib
ERK VX-11e
FGFR PD-173074
BGJ398
LCK A770041
mTOR RAD001
p53 activation PRIMA-1MET
PI3K Alpelisib
Pictilisib
PI3K, mTORC GSK2126458
NVP-BEZ235
RAF RAF265
vemurafenib
AZ628
Dabrafenib
Primary Target (group)
Drug Name
Trametinib
ABL Nilotinib
ABL, SRC Dasatinib
ATP synthase Oligomycin
BRD I-BET
JQ1
ERBB Afatinib
Erlotinib
Lapatinib
Neratinib
ERK VX-11e
IGF11R, InsR BMS-754807
IGF1R Linsitinib
LCK A770041
PI3K Alpelisib
Pictilisib
PI3K, mTORC GSK2126458
NVP-BEZ235
RAF RAF265
vemurafenib
AZ628
Dabrafenib
mTOR RAD001
mTORC OSI-027
AZD8055


E

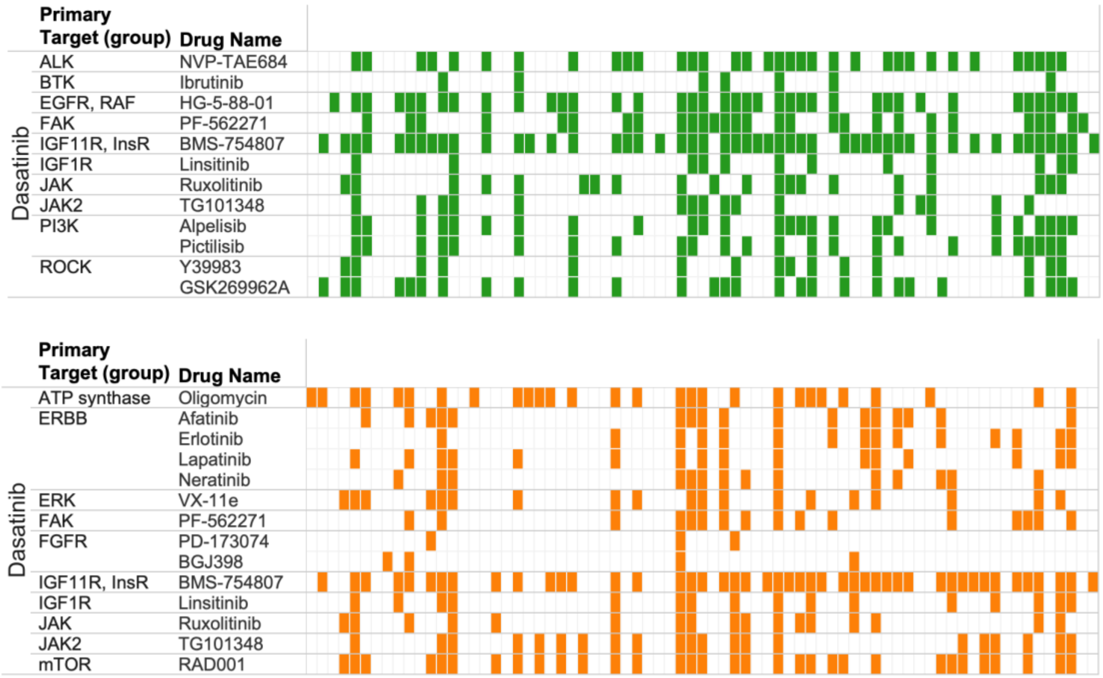

Primary Target (group)
Drug Name
Dasatinib
ALK NVP-TAE684
BTK Ibrutinib
EGFR, RAF HG-5-88-01
FAK PF-562271
IGF11R, InsR BMS-754807
IGF1R Linsitinib
JAK Ruxolitinib
JAK2 TG101348
PI3K Alpelisib
Pictilisib
ROCK Y39983
GSK269962A
Primary Target (group)
Drug Name
Dasatinib
ATP synthase Oligomycin
ERBB Afatinib
Erlotinib
Lapatinib
Neratinib
ERK VX-11e
FAK PF-562271
FGFR PD-173074
BGJ398
IGF11R, InsR BMS-754807
IGF1R Linsitinib
JAK Ruxolitinib
JAK2 TG101348
mTOR RAD001

**Supp Figure 3:** Pattern of synergies and HSA events across related drugs. Several examples of patterns obtained for drugs with related targets are shown for some of the anchors. A-E: Cell lines (columns) are in the same order across rows to allow for pattern comparison. Cell lines harboring HSA (orange) or synergy (green) are colored accordingly. The same threshold for synergy and HSA are used across all panels. Anchors for the different panels are: A, Alpelisib (PI3Kalpha); B, Pictilisib (pan PI3K); C, Palbociclib (CDK4/6); D, Trametinib (MEK1/2); E, Dasatinib (ABL, SRC).

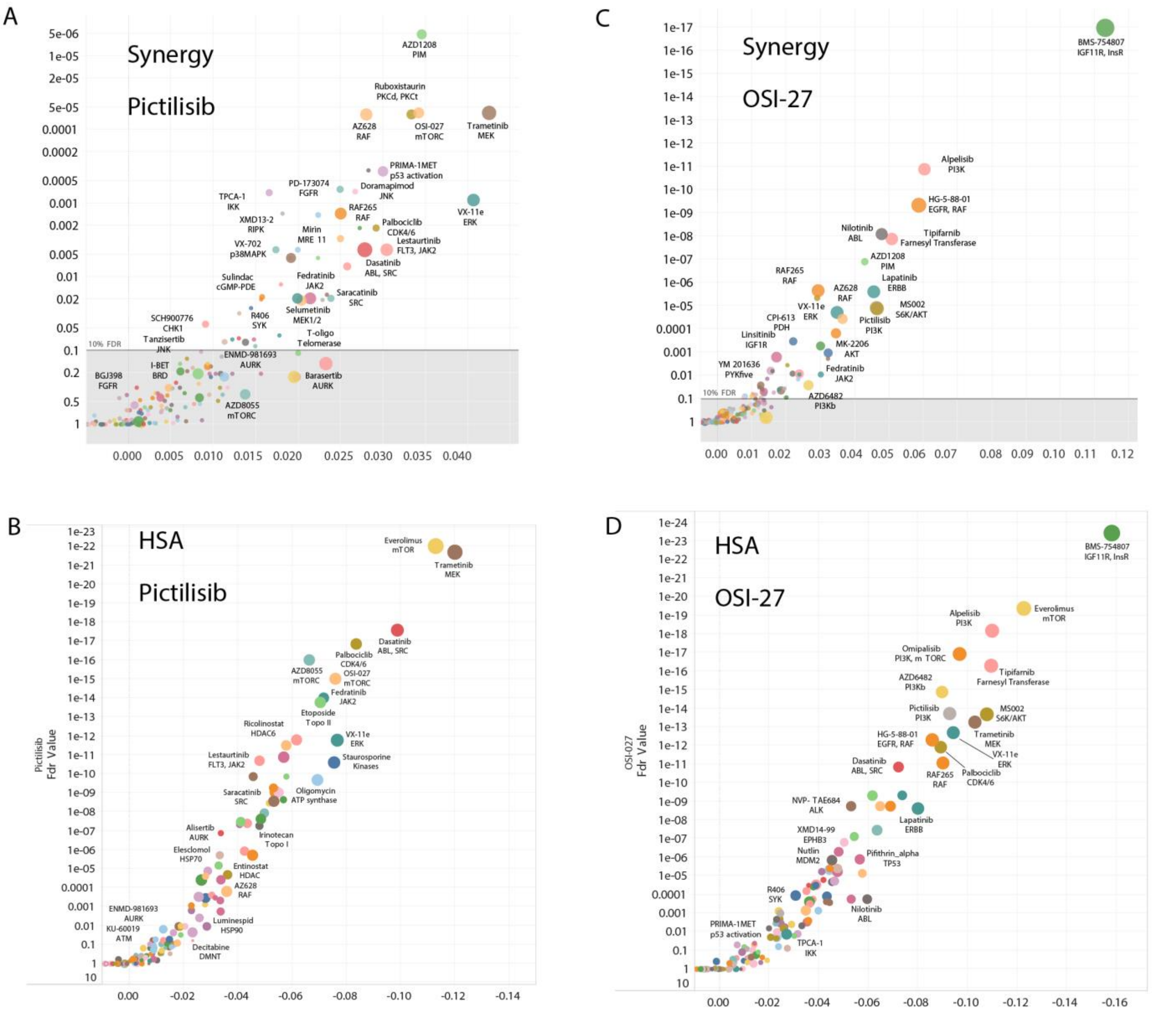

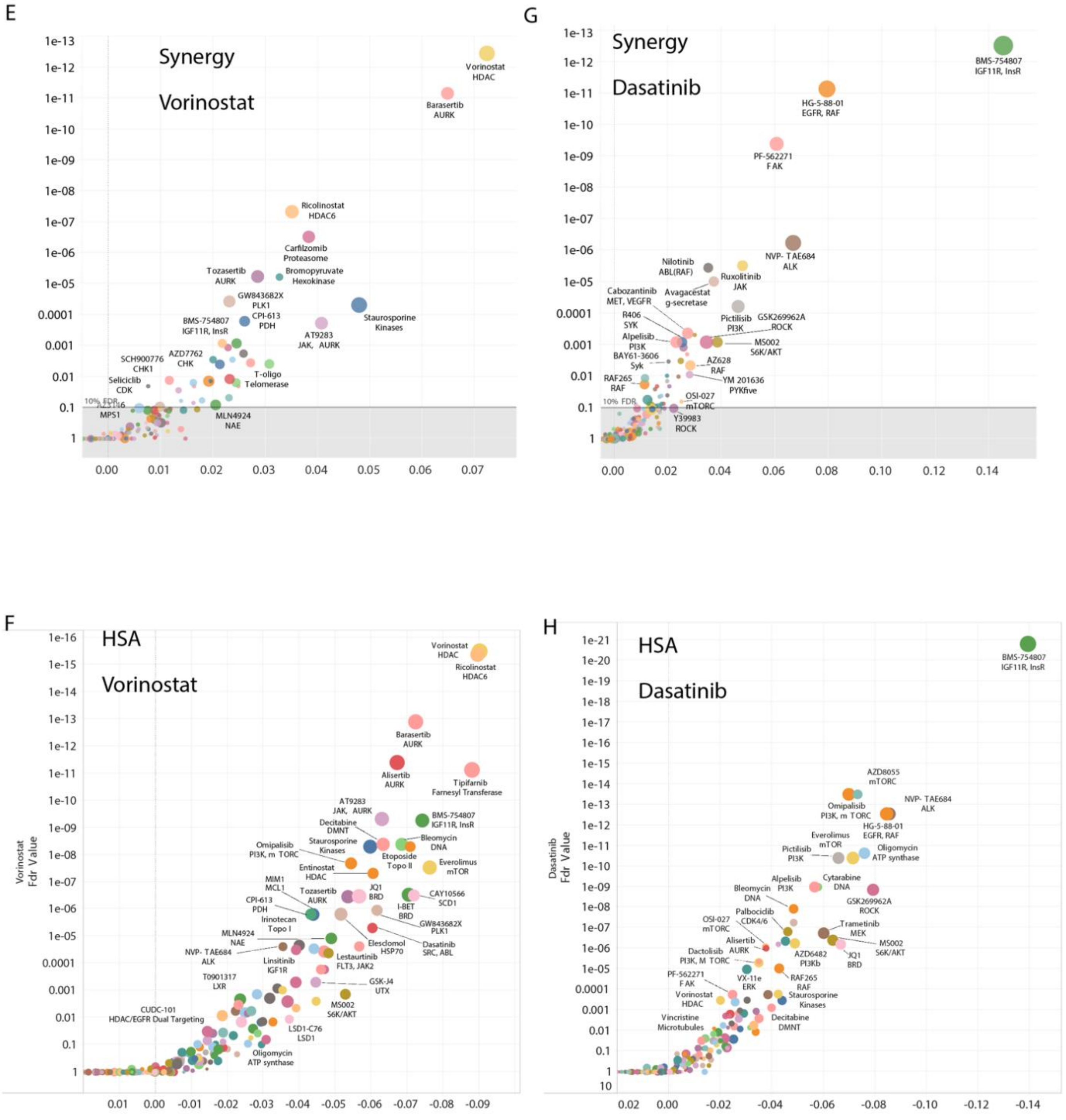


**Supp Figure 4:** Distribution of synergy and HSA scores for selected anchors illustrates global consistency of the two scoring metrics. The impact score for each combination is plotted. X axis: Differential median synergy score across cell lines corresponding to the Log10 ratio of the median synergy (or HSA) score for the indicated library drug over median score of all other drugs. Y axis: FDR value for statistical enrichment of synergies for the plotted combination over all other tested combinations (with the same anchor). The size of the dots represent the percentile of synergy scores for a given combination falling within the top 5% of all synergy scores for the whole screen (all anchors). Anchors: Pictilisib (pan PI3K), OSI-27 (mTORC), Votinostat (HDAC), Dasatinib (ABL, SRC).

A

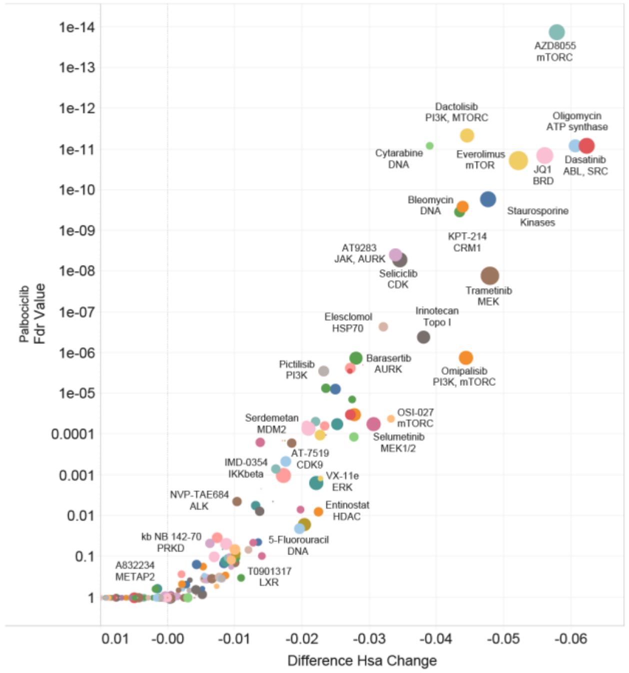

Palbociclib
Fdr Value
1e-14
1e-13
1e-12
1e-11
1e-10
1e-09
1e-08
1e-07
1e-06
1e-05
0.0001
0.001
0.01
0.1
1
0.01
-0.00
-0.01
-0.02
-0.03
-0.04
-0.05
-0.06
Difference Hsa Change
AZD8055
mTORC
Dactolisib
PI3K, MTORC
Oligomycin
ATP synthase
Cytarabine
DNA
Everolimus
mTOR
JQ1
BRD
Dasatinib
ABL, SRC
Bleomycin
DNA
Staurosporine
Kinases
KPT-214
CRM1
AT9283
JAK, AURK
Seliciclib
CDK
Trametinib
MEK
Elesclomol
HSP70
Irinotecan
Topo I
Pictilisib
PI3K
Barasertib
AURK
Omipalisib
PI3K, mTORC
Serdemetan
MDM2
OSI-027
mTORC
Selumetinib
MEK1/2
AT-7519
CDK9
IMD-0354
IKKbeta
VX-11e
ERK
NVP-TAE684
ALK
Entinostat
HDAC
kb NB 142-70
PRKD
5-Fluorouracil
DNA
A832234
METAP2
T0901317
LXR


B

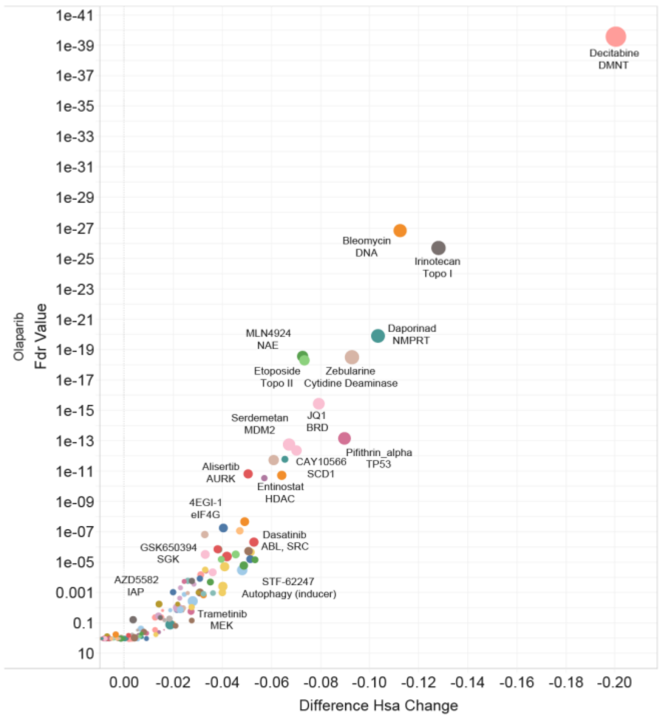

Olaparib
Fdr Value
1e-41
1e-39
1e-37
1e-35
1e-33
1e-31
1e-29
1e-27
1e-25
1e-23
1e-21
1e-19
1e-17
1e-15
1e-13
1e-11
1e-09
1e-07
1e-05
0.001
0.1
10
0.00
-0.02
-0.04
-0.06
-0.08
-0.10
-0.12
-0.14
-0.16
-0.18
-0.20
Difference Hsa Change
Decitabine
DMNT
Bleomycin
DNA
Irinotecan
Topo I
MLN4924
NAE
Daporinad
NMPRT
Etoposide
Topo II
Zebularine
Cytidine Deaminase
Serdemetan
MDM2
JQ1
BRD
Pifithrin_alpha
TP53
CAY10566
SCD1
Alisertib
AURK
Entinostat
HDAC
4EGI-1
eIF4G
Dasatinib
ABL, SRC
GSK650394
SGK
STF-62247
Autophagy (inducer)
AZD5582
IAP
Trametinib
MEK


C

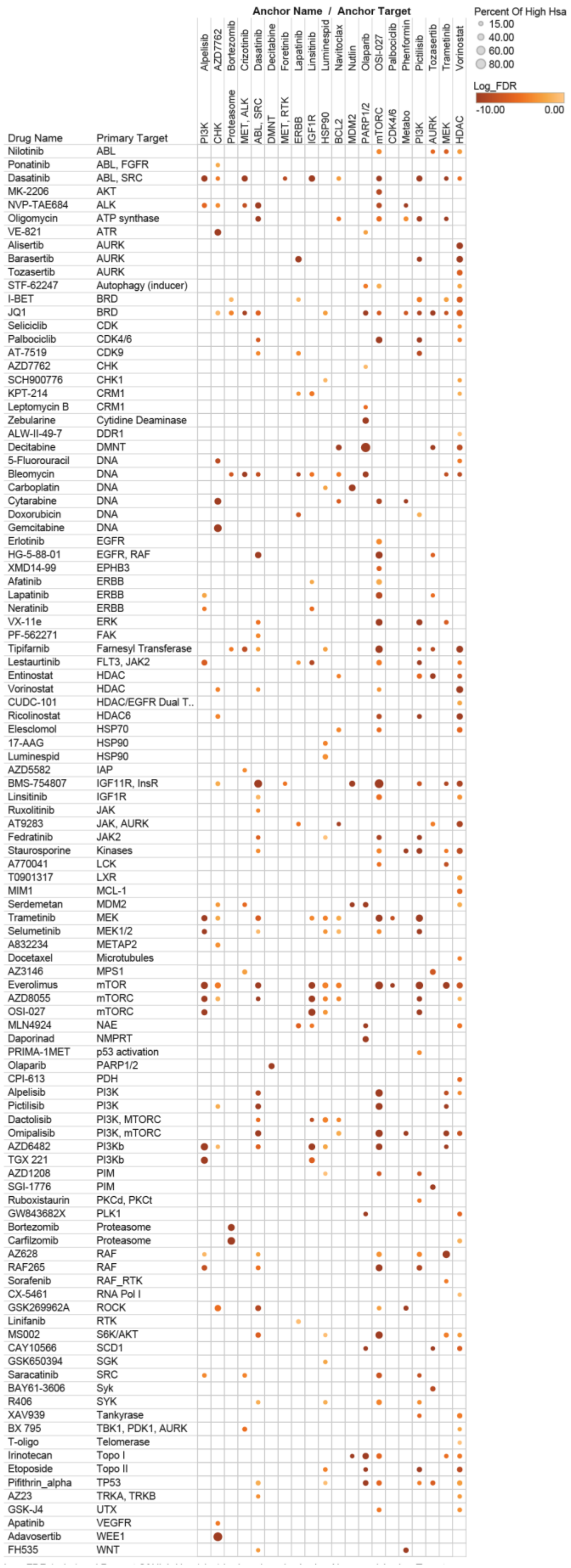

Anchor Name / Anchor Target
Alpelisib
AZD7762
Bortezomib
Crizotinib
Dasatinib
Decitabine
Foretinib
Lapatinib
Linsitinib
Luminespid
Navitoclax
Nutlin
Olaparib
OSI-027
Palbociclib
Phenformin
Pictilisib
Tozasertib
Trametinib
Vorinostat
PI3K
CHK
Proteasome
MET, ALK
ABL, SRC
DMNT
MET, RTK
ERBB
IGF1R
HSP90
BCL2
MDM2
PARP1/2
mTORC
CDK4/6
Metabo
PI3K
AURK
MEK
HDAC
Percent Of High Hsa
15.00
40.00
60.00
80.00
Log_FDR
-10.00
0.00
Drug Name
Primary Target
Nilotinib
ABL
Ponatinib
ABL, FGFR
Dasatinib
ABL, SRC
MK-2206
AKT
NVP-TAE684
ALK
Oligomycin
ATP synthase
VE-821
ATR
Alisertib
AURK
Barasertib
AURK
Tozasertib
AURK
STF-62247
Autophagy (inducer)
I-BET
BRD
JQ1
BRD
Seliciclib
CDK
Palbociclib
CDK4/6
AT-7519
CDK9
AZD7762
CHK
SCH900776
CHK1
KPT-214
CRM1
Leptomycin B
CRM1
Zebularine
Cytidine Deaminase
ALW-II-49-7
DDR1
Decitabine
DMNT
5-Fluorouracil
DNA
Bleomycin
DNA
Carboplatin
DNA
Cytarabine
DNA
Doxorubicin
DNA
Gemcitabine
DNA
Erlotinib
EGFR
HG-5-88-01
EGFR, RAF
XMD14-99
EPHB3
Afatinib
ERBB
Lapatinib
ERBB
Neratinib
ERBB
VX-11e
ERK
PF-562271
FAK
Tipifarnib
Farnesyl Transferase
Lestaurtinib
FLT3, JAK2
Entinostat
HDAC
Vorinostat
HDAC
CUDC-101
HDAC/EGFR Dual T..
Ricolinostat
HDAC6
Elesclomol
HSP70
17-AAG
HSP90
Luminespid
HSP90
AZD5582
IAP
BMS-754807
IGF11R, InsR
Linsitinib
IGF1R
Ruxolitinib
JAK
AT9283
JAK, AURK
Fedratinib
JAK2
Staurosporine
Kinases
A770041
LCK
T0901317
LXR
MIM1
MCL-1
Serdemetan
MDM2
Trametinib
MEK
Selumetinib
MEK1/2
A832234
METAP2
Docetaxel
Microtubules
AZ3146
MPS1
Everolimus
mTOR
AZD8055
mTORC
OSI-027
mTORC
MLN4924
NAE
Daporinad
NMPRT
PRIMA-1MET
p53 activation
Olaparib
PARP1/2
CPI-613
PDH
Alpelisib
PI3K
Pictilisib
PI3K
Dactolisib
PI3K, MTORC
Omipalisib
PI3K, mTORC
AZD6482
PI3Kb
TGX 221
PI3Kb
AZD1208
PIM
SGI-1776
PIM
Ruboxistaurin
PKCd, PKCt
GW843682X
PLK1
Bortezomib
Proteasome
Carfilzomib
Proteasome
AZ628
RAF
RAF265
RAF
Sorafenib
RAF_RTK
CX-5461
RNA Pol I
GSK269962A
ROCK
Linifanib
RTK
MS002
S6K/AKT
CAY10566
SCD1
GSK650394
SGK
Saracatinib
SRC
BAY61-3606
Syk
R406
SYK
XAV939
Tankyrase
BX 795
TBK1, PDK1, AURK
T-oligo
Telomerase
Irinotecan
Topo I
Etoposide
Topo II
Pifithrin_alpha
TP53
AZ23
TRKA, TRKB
GSK-J4
UTX
Apatinib
VEGFR
Adavosertib
WEE1
FH535
WNT

**Supp Figure 5:** HSA Scores for selected drugs and overall HSA score matrix. A and B, HSA impact scores for the anchors palbociclib (CDK4/6) and Olaparib (PARP). C: Overview of HSA scores across tested combination. Combinations yielding HSA in at least 15% of the cell lines tested are shown. The size of the dot corresponds to the percentile of cell lines with HSA and the shade to the statistical enrichment of HSA events for the indicated combination over all other tested combinations (darker corresponds to lower FDR).

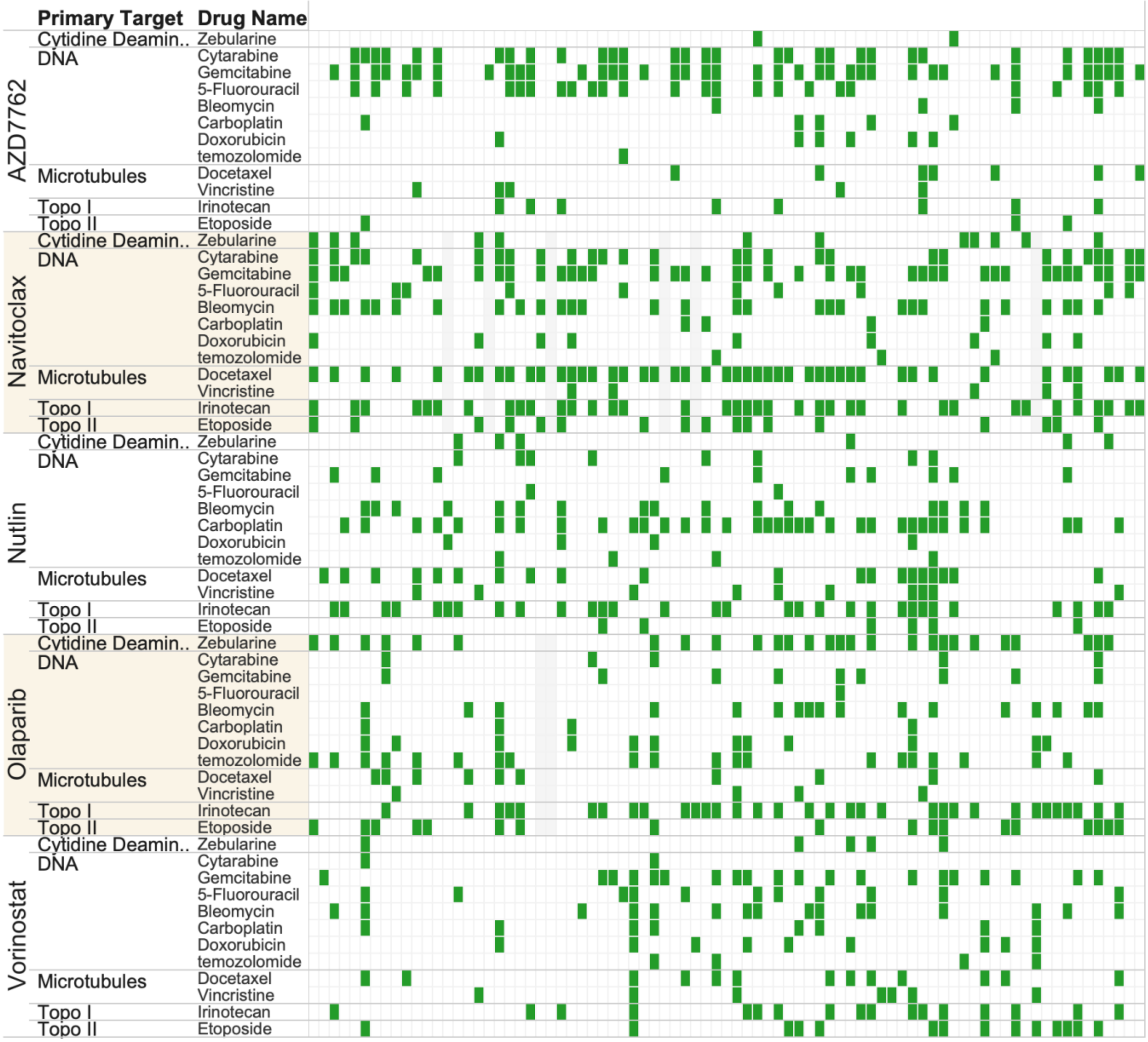

**Supp Figure 6:** Pattern of synergy score for cytotoxic drugs. For each of the indicated anchor drugs (vertical labels) the pattern of synergy for a given library drug (horizontal label) is shown by a color box when the chosen threshold of 0.8 synergy score is passed. Cell lines are in same order across all rows. Gray shading corresponds to missing data.

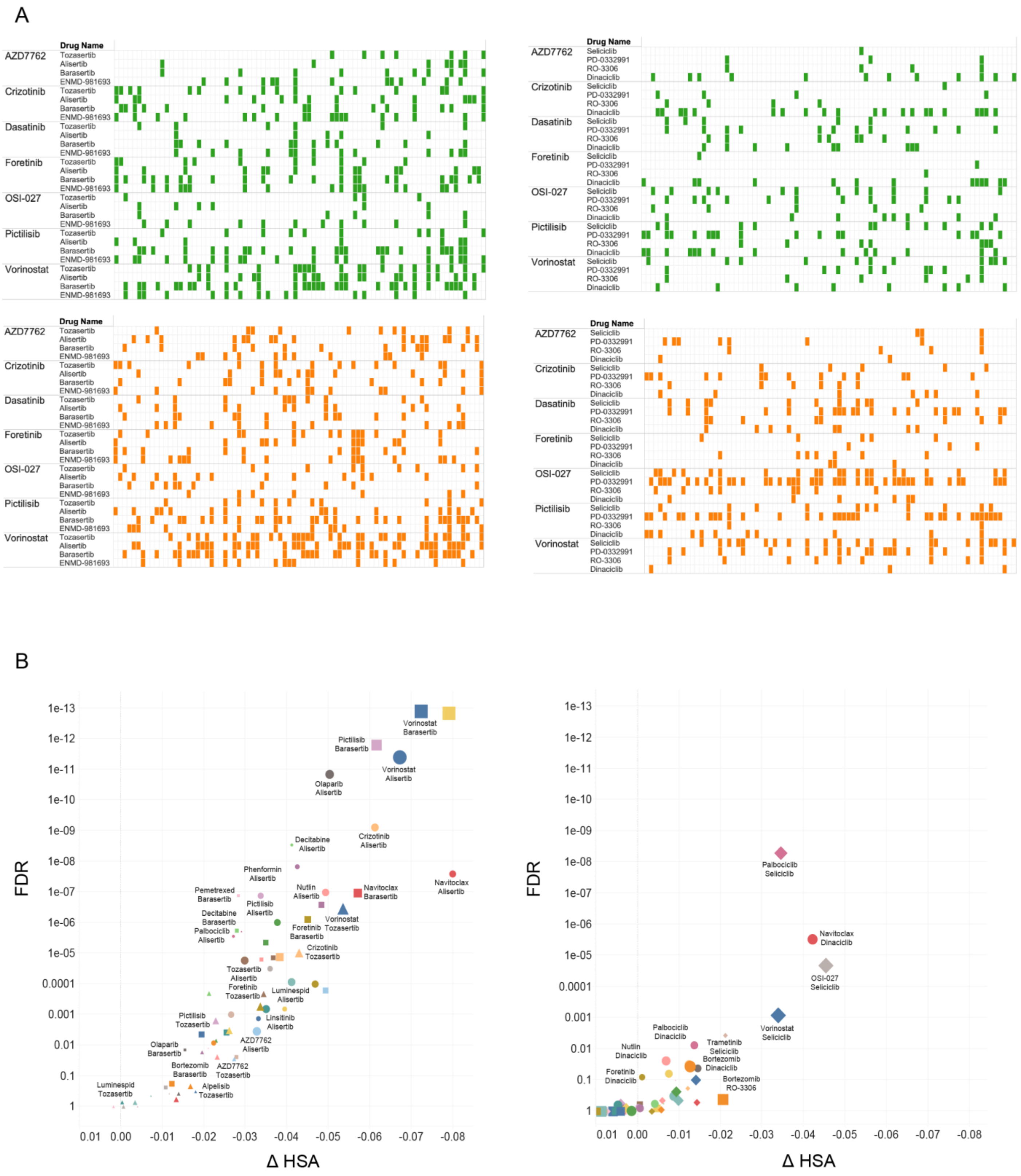

**Supp Figure 7**: Aurora kinase inhibitors are engaged more frequently in synergistic and HSA events than CDK inhibitors. A, Pattern of synergy and HSA events across selected anchors for the library drugs targeting Aurora Kinases and CDKs. B, Impact score graphs for Aurora Kinase and CDK inhibitors.

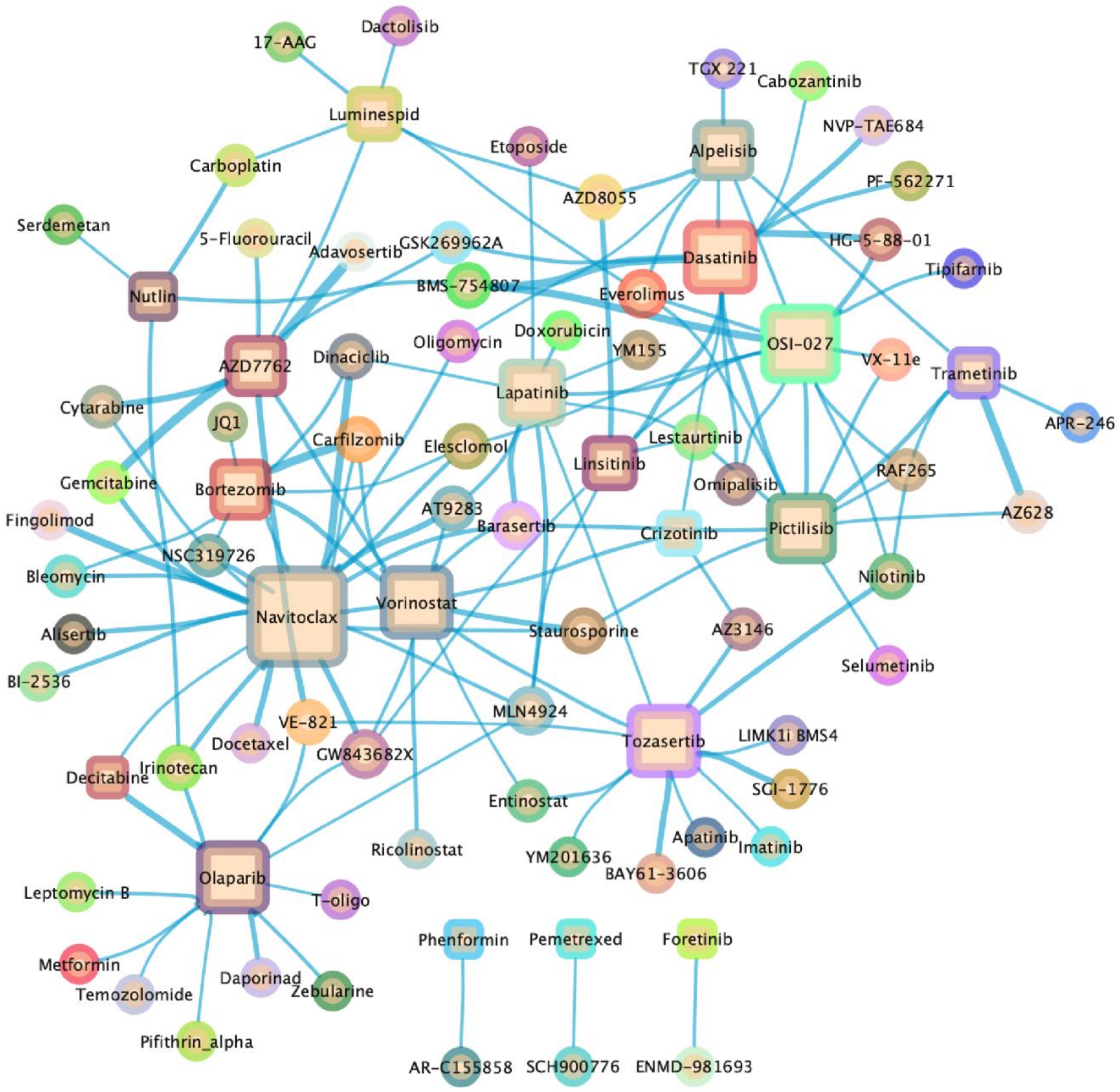


**Supp Figure 8:** Network representation of drug-drug interactions for top 5% synergistic events across all drugs and anchors. Anchor drugs are represented as squares and library drugs as circles. The size of the anchor nodes is proportional to the number of synergistic events the corresponding drug is involved in. The thickness of the edge between two drugs corresponds to the median value of the synergy scores across all cell lines.

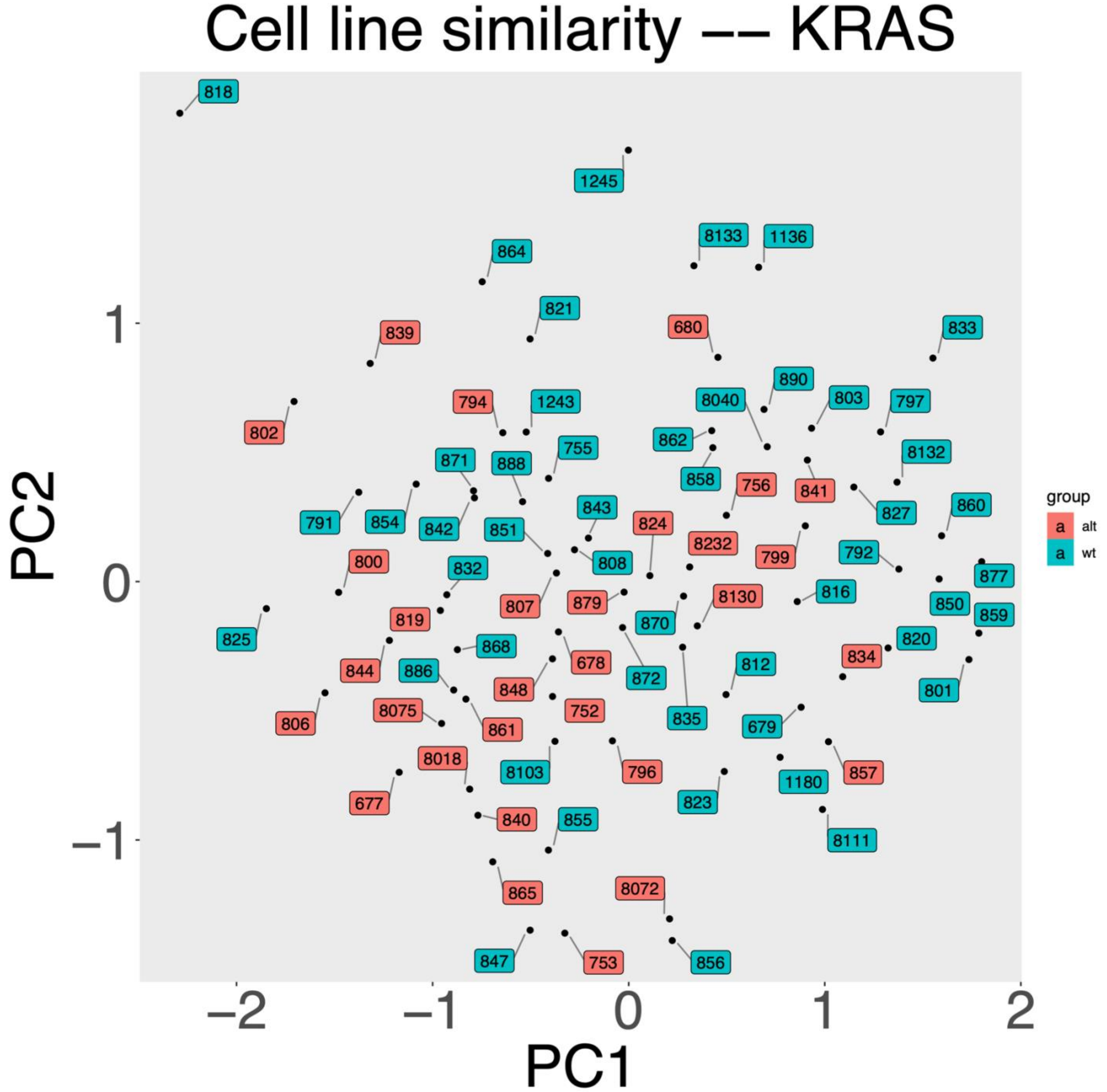


**Supp Figure 9:** Analysis of synergy patterns across drugs for each cell line does not differentiate KRAS WT from KRAS mutant cell lines. Principle component analysis was used to analyze the proximity of cell lines based on the pattern of synergistic events across all anchors and library drugs. Cell lines are color coded based on KRAS mutational status.
